# Bradykinin Contributes to Vasogenic Edema in Murine Experimental Cerebral Malaria

**DOI:** 10.64898/2026.02.23.704410

**Authors:** Alessandro de Sa Pinheiro, Douglas E. Teixeira, Rodrigo P. Silva-Aguiar, Young Jun Shim, Alona A. Merkulova, Sadiq Silbak, Yelenna Skomorovska-Prokvolit, David Midem, Sidney Ogolla, Bjoern B. Burckhardt, Tanja Gangnus, Julio Scharfstein, Celso Caruso-Neves, Owen J.T. McCarty, David Gailani, Michael Bader, Philip J. Rosenthal, Arlene E. Dent, Chris J. Janse, Keith R. McCrae, Ana Acacia de Sa Pinheiro, James W. Kazura, Alvin H. Schmaier

## Abstract

Cerebral malaria (CM) due to *Plasmodium falciparum* (*Pf*) infection is a major cause of death in African children. Bradykinin (BK) is a mediator of vasogenic edema. It could contribute to the pathogenesis of central nervous system malaria in Kenyan children and *P. berghei* ANKA (*PbA*) infected C57BL/6J mice with experimental cerebral malaria. Cleaved plasma high molecular weight kininogen (cHK) is a marker for prior BK release. 40% of children with central nervous system malaria had plasma cHK versus 18% of children with uncomplicated malaria. Wild-type *PbA*-infected mice had circulating plasma cHK, elevated BK levels, and reduced HK and prekallikrein levels. HK null (*Kng1^-/-^*), combined BK B1 and B2 receptor null (*Bdkrb1^-/-^ / Bdkrb2^-/-^*), BK B2 (*Bdkrb2^-/-^*) or BK B1 (*Bdkrb1^-/-^*) receptor null mice were protected from neurologic deterioration and brain edema compared to wild-type mice. *F12^-/-^*mice were not protected from neurological deterioration.

Prekallikrein null (*Klkb1^-/-^*), prolylcarboxypeptidase hypomorphs (*Prcp^gt/gt^*), and brain endothelial cell conditional knockout of PRCP (*Prcp^fl/fl^* Cre) mice had reduced neurologic deterioration and brain edema. Adjuvant plasma kallikrein inhibition combined with artesunate treatment of *PbA*-infected mice reversed neurologic deterioration and brain edema and prolonged survival relative to artesunate alone. BK-induced vasogenic edema contributes to human and murine CM.

## Introduction

Malaria is a major cause of death in children worldwide. According to the World Health Organization (WHO), there were an estimated 263 million malaria cases leading to 597,000 deaths in 2023 (1).

Seventy-six percent of these deaths occurred in African children under 5 years old. Children with cerebral malaria (CM), a severe malaria phenotype, can die despite artesunate therapy (2).

Moreover, 30% of pediatric CM survivors experience post-recovery neurocognitive, motor, and learning deficits that have unclear significance to future growth and development (3–5).

Studies of pediatric CM in endemic sites by magnetic resonance imaging show that brain swelling is associated with vasogenic edema (breakdown of the blood brain barrier) and obstructed cerebral venous blood outflow (6,7). In fatal CM, progressive brain swelling leads to brainstem dysfunction and uncal herniation. Features of pediatric CM pathogenesis include a high *Plasmodium falciparum* (*Pf*) biomass with circulating DNA accompanied by sequestered parasitized erythrocytes in cerebral microvasculature, endothelial cell activation, and neuro-inflammation with neutrophil and CD4 and CD8 T cell infiltration (8–13). The proximal mechanism(s) that initiate and mediate progressive brain swelling is/are not known. An investigation by Higgins *et al*. showed that administration of angiopoietin 1 in conjunction with artesunate increased survival of mice with experimental cerebral malaria (ECM) (14). This ground-breaking study shows that maximizing tight junctions of the blood brain barrier with angiopoietin 1 in conjuction with anti-parasite therapy reduces vasogenic edema and improves survival. The lack of adjuvant treatments to reduce brain vasogenic edema in children with CM is a major unmet need.

Previously our group demonstrated that *Pf* culture medium co-incubated with human brain microvascular endothelial cells induces disruption of intercellular junctions through BK B2 and B1 receptors (17). Other investigations suggest that BK and related plasma proteins contribute to malarial disease pathogenesis (18). BK is a well-established mediator of increased vascular permeability and initiates edema formation in hereditary angioedema (HAE). We hypothesize that similar mechanisms underlie endothelial barrier dysfunction and vasogenic edema in CM. Types I and II HAE are disorders characterized by acute bouts of localized tissue edema due to C1 Inhibitor (C1INH) deficiency or defects. This disorder is initiated by activation of plasma prekallikrein (PK) to plasma kallikrein (PKa) that proteolyzes high molecular weight kininogen (HK) with release of BK, leaving residual circulating plasma cleaved HK (cHK) (17–22). BK is a short-lived 9 amino acid peptide generated by proteolytic cleavage of HK and low molecular weight kininogen, both encoded by the single *KNG1* gene. Measurement of BK levels *in viv*o is challenging because the bioactive peptide is rapidly degraded and cleared from plasma with a half-life of ∼34 sec (23,24). Cleaved HK, on the other hand, has a half-life of 10 h and serves as an established biomarker of prior BK generation in plasma (20–22, 25). In this study, we examined plasma samples for detection of cHK and HK levels from children who presented with central nervous system malaria diagnosed at a rural hospital in western Kenya. The median (interquartile) Blantyre coma score was 2 (1–3) (26). The term central nervous system malaria (CNS-M), not CM, is used in describing these patients since hospital personnel did not have the capacity to perform fundoscopic examination for malaria retinopathy (27). Using the ECM model of *Plasmodium berghei* ANKA (*PbA*)-infected C57BL/6J mice, we examined plasma for activation of factor XII (FXII) and the kallikrein/kinin system (KKS). We also examined the mouse genetic deletion models for FXII and the proteins of the KKS and used inhibitors of select proteins of this system to determine their influence on neurologic deterioration, vasogenic brain edema, and survival in ECM.

## Results

*Changes in plasma HK levels in Kenyan children with central nervous system malaria (CNS-M), uncomplicated malaria (UM), acute febrile non-malaria illness (AFN) or healthy community controls (Healthy).* Plasma from venous blood collected in heparin anticoagulant was isolated from Kenyan children with CNS-M, UM, AFN, and healthy community controls. Demographic and clinical characteristics of these children are presented in **Table** 1 (26). Children with CNS-M were younger than those with UM. The Blantyre coma scores of the CNS-M children were < 3. Plasma *Pf*HRP2 levels, an estimate of the total *Pf* biomass, were significantly higher in children with CNS-M than those with UM (**Table 1**).

**Table 1.** Clinical Characteristics of Kenyan Children with Malaria and Controls at Entry.

|  | CNS-M<sup>1</sup> | UM<sup>1</sup> | AFN<sup>1</sup> | Healthy<sup>1</sup> |
| --- | --- | --- | --- | --- |
| Number | 20 | 17 | 12 | 15 |
| Age (years)<sup>2</sup> | 2.8 (2.3, 4.4) | 5.2 (4.1, 8.4) | 5.3 (4.1, 7.9) | 5.3 (3.7, 7.3) |
| Temperature | >37.8°C | >37.8°C | >37.8°C | ≤37.8°C |
| Blantyre score | 0-3 | 5 | 5 | 5 |
| Blood smear for *Pf* | positive | positive | negative | negative |
| *Pf*HRP2 (ng/ml)<sup>2,3</sup> | 486 (137, 1396) | 43 (14,166) | 0 | 0 |
| Hgb (g/dl)<sup>2,3</sup> | 10.6 (9.4,11.41) | 11.1 (9.7,13.5) | 10.4 (7.5, 15) | - |

<sup>1</sup> CNS-M = central nervous system malaria; UM = uncomplicated malaria; AFN = acute febrile non-malaria illness; and Healthy = healthy community control children
<sup>2</sup> median age and interquartile range (IQR)
<sup>3</sup> Comparison of CNS-M vs UM values show a P = 0.0005, Mann-Whitney two-tailed t test.
*Pf*HRP2 is *Plasmodium falciparum* histidine-rich protein 2 and Hgb is hemoglobin.

Plasma HK levels were measured by a competitive ELISA using a rabbit polyclonal antibody that detected total HK light chain protein, i.e. both intact HK and cHK (**Fig. 1A**). CNS-M patients at hospital entry (CNS-M-En) had HK levels that were significantly lower (P = 0.03) than patients with UM. UM patients had HK levels significantly higher than those with AFN (P = 0.003) and Healthy (P < 0.0001) children who were blood smear negative for Plasmodium parasites and PCR negative for *Pf* 18s ribosomal gene. Plasma HK of CNS-M patients at hospital discharge (CNS-M-D) following artesunate treatment and clinical recovery were significantly increased (P = 0.02) relative to CNS-M-En (**Fig. 1B**). CNS-M values at hospital discharge were like those of children with UM and significantly higher than the AFN (P = 0.01) and Healthy (P < 0.0001) children (**Table 2**).

**Figure 1.**
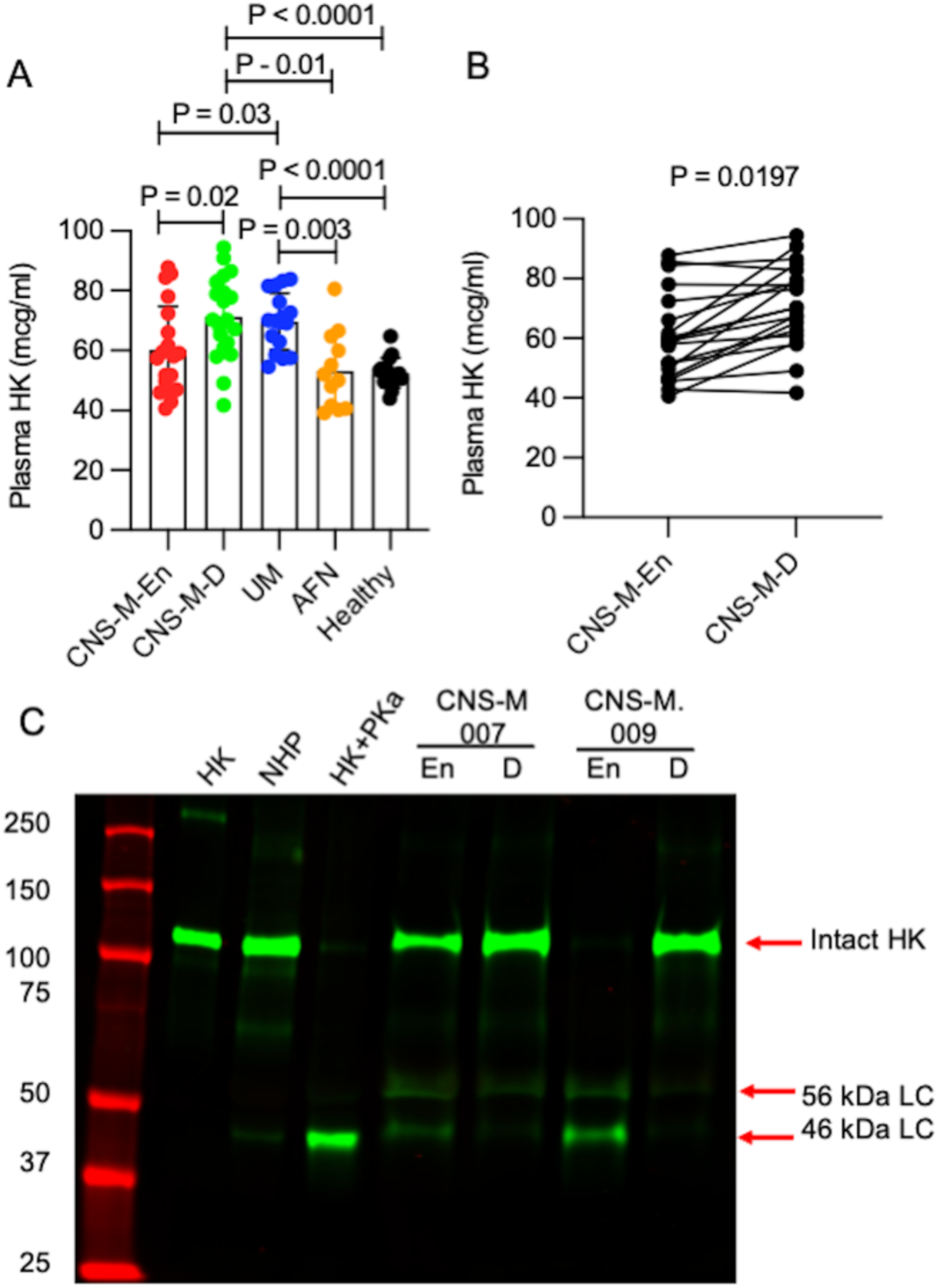
Investigations on plasma high molecular weight kininogen (HK) on Kenyan children’s samples. Panel A. Plasma HK levels in children with central nervous system malaria (CNS-M) at hospital entry (CNS--En, n =20), at hospital discharge (CNS-M-D, n = 20), uncomplicated malaria (UM, n = 17), acute febrile non-malaria illness (AFN, n = 12), and healthy community control children (Healthy, n =15). A goat-anti-human antibody reared with purified HK and absorbed with Fitzgerald plasma was used for the CELISA. All values shown are plasma HK levels in mcg/ml (mean ± SD). Comparisons shown are for all significant difference between groups. The absence of comparison between groups indicates non-significant differences. Non-significant differences between groups were between CNS-M-D and UM, AFN and Healthy, and CNS-M-En and AFN or Healthy. The statistical analysis between groups was performed by a two-tailed t test, P < 0.05 was considered significant. Panel B. Paired samples of plasma HK concentrations of CNS-M patients at hospital entry (CNS-M-En) and discharge (CNS-M-D) by group paired two-tailed t-test. Panel C. Immunoblot with reduced samples on a 7.5% SDS-PAGE of two CNS-M patient’s plasma (CNS-M-007, CNS-M-009) at hospital entry (En) and discharge (D) using a rabbit antibody to a human HK D5 peptide (anti-LDDDLEHQGGHVLDHGHKHK-HGHGHGKHKNKGKK-NGK). On the immunoblot, HK represents pure protein, NHP represents an immunoblot of pooled normal human plasma HK, and HK+PKa is an immunoblot of a purified human plasma kallikrein cleavage of pure human HK. The two arrows on the right side of the immunoblot point out the 56 and 46 kDa light chains (LC) of cleaved HK (cHK). Note all plasmas (normal pooled or patient samples) were added to the gel as 5 μl of a 1:10 dilution of plasma.

**Table 2.** Plasma High Molecular Weight Kininogen in Malaria Samples.

| Sample<sup>1</sup> | Number | HK concentration (mcg/ml)<sup>2</sup> |
| --- | --- | --- |
| CNS-M-En | 20 | 60 ± 15 |
| CNS-M-D | 20 | 71 ± 14 |
| UM | 17 | 70 ± 10 |
| AFN | 12 | 53 ± 13 |
| Healthy | 15 | 53 ± 5 |

<sup>1</sup> Children plasma samples with central nervous system malaria are: CNS-M-En = CNS-M at entry; CNS-M-D = CM at discharge; UM = UM; AFN = Acute febrile non-malaria illness; and Healthy = healthy community control children.
<sup>2</sup> Each value represents the mean ± 1 SD.

Since differences in the quantity of total HK measured were not strikingly different among the patient groups, immunoblot studies for plasma HK were performed to see if any 56 and 46 kDa cHK, a validated marker of BK generation, was present (19–22). These investigations showed that 8 out of 20 CNS-M-En (40%) plasma samples collected before starting artesunate treatment had circulating cHK. Four of these 8 children had Blantyre scores of < 2.0; 4 children had Blantyre scores of 3. Using an antibody to domain 5 of human HK (**Fig. 1C)**, pure intact HK (lane HK) and HK in normal human plasma (lane NHP) on reduced SDS-PAGE were single ∼120 kDa bands. When the purified HK was treated with purified plasma kallikrein (HK+PKa), intact HK was mostly cleaved to a terminal 42-46 kDa light chain. These samples were used as controls for immunoblot studies of plasma from CNS-M patients. In patient CNS-M 007, intact HK at 120 kDa was reduced at hospital entry vs. discharge with intermediate 56 kDa and terminal 46 kDa bands of the cleaved light chain of HK (two arrows on the right of the blot). At the time of hospital discharge, plasma from patient CNS-M 007 had mostly intact HK with a faint 56 kDa light chain remaining. In **Fig. 1C**, patient CNS-M 009 had no intact 120 kDa HK at study entry; only the two HK light chain fragments of cHK were detected at 56 and 46 kDa. At hospital discharge, plasma from this patient only contained intact 120 kDa HK. In contrast to the CNS-M patients, only 3 of 17 (18%) of UM patients had circulating cHK at study entry. None of the plasma immunoblot samples from children with AFN or healthy uninfected children had cHK. Thus, children with CNS-M and to less extent, UM, were associated with the presence of cHK, a structure consistent with prior *in vivo* BK release (19–22).

*Hematologic and coagulation parameters in C57BL/6J mice with ECM-.* Functional assays of the kallikrein/kinin system could not reliably be performed on the Kenyan children blood samples because they were collected in heparin. We therefore used the ECM mouse model to evaluate the hematologic and coagulation parameters associated with malaria pathogenesis. *PbA* murine malaria is used because *P. falciparum* only infects *Homo sapiens*. After intraperitoneal (IP) inoculation with 10^6^ *PbA*-infected red blood cells (iRBCs) (**Fig. 2**), murine blood was collected at day 5 post-infection at the time of onset of neurologic deterioration measured by the SHIRPA score (28). Relative to uninfected control mice, *PbA*-infected mice had significantly lower (P < 0.0001) white blood cell counts, lymphocyte counts, hematocrits, RBC mean corpuscular volume, and platelet counts. The mean corpuscular hemoglobin concentration in the infected mice was significantly higher than uninfected mice. The mean activated partial thromboplastin time (aPTT) (P = 0.001) and prothrombin time (PT) (P = 0.008) were prolonged in *PbA*-infected mice compared to uninfected mice. There was a trimodal distribution of the PT that included a cluster five mice with markedly prolonged PTs, a group with moderately prolonged PTs, and a third group with PTs comparable to uninfected mice. The clinical significance of these prothrombin time findings is presently not known since these three groups of mice had the same advanced, objective neurologic deficits on the SHIRPA score at the time of euthanasia on day 5 post-infection.

**Figure 2.**
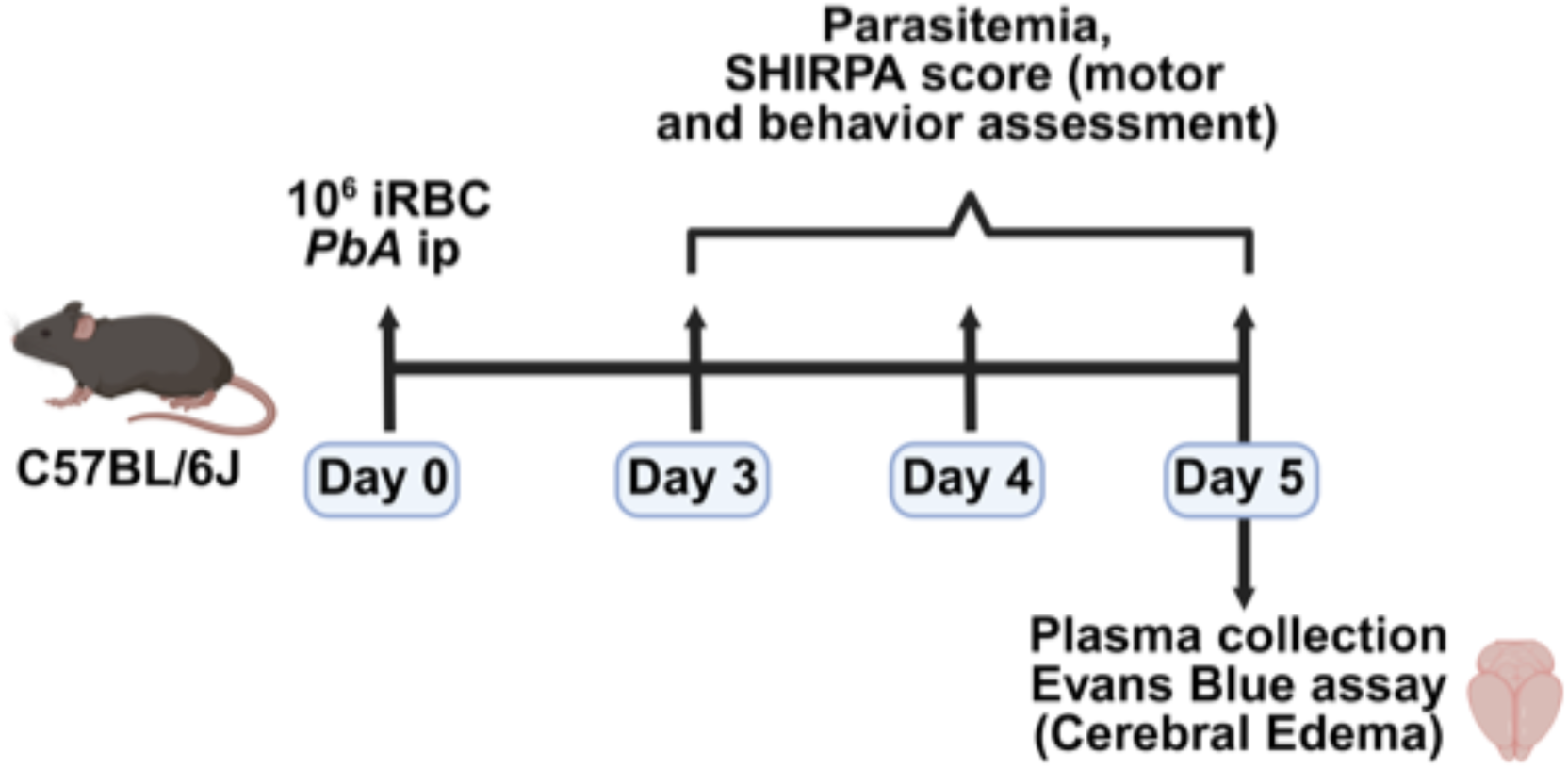
Murine experimental cerebral malaria model based on *Plasmodium berghei* ANKA (*PbA*). On day 0, the mouse was injected i.p. with 10^6^ *P. berghei* ANKA iRBCs in 100 μl buffer. On days 3, 4 and 5 after infection, parasitemia was measured and the animal underwent a detailed SHIRPA score assessment of neurologic activity. On day 5, when wild type C57BL/6J mice demonstrated overt neurologic deterioration, mice had blood collected for parasitemia and received Evans Blue dye injection. The animals then were euthanized, and brains were excised for examination.

We next examined if *PbA*-infected mice had changes in the levels of the proteins of the plasma kallikrein/kinin system and factor XII. Plasma HK levels in *PbA*-infected mice (0.78 ± 0.08 IU/ml coagulant activity and 0.59 ± 0.09 IU/ml HK antigen) were lower than values in uninfected mice (1.2 ± 0.07 IU/ml and 1.0±0.06 IU/ml, P = 0.016 and P = 0.0024, respectively) (**Fig. 3A**). An immunoblot of mouse plasma using an antibody to murine HK domain 6 (D6) revealed two bands: one at ∼120 kDa and a second at 100 kDa (ΔmHK-D5) in both uninfected and infected mice (**Fig. 3B**). This latter form of murine HK had been previously described and corresponded to HK lacking domain 5 (D5) (29).

**Figure 3.**
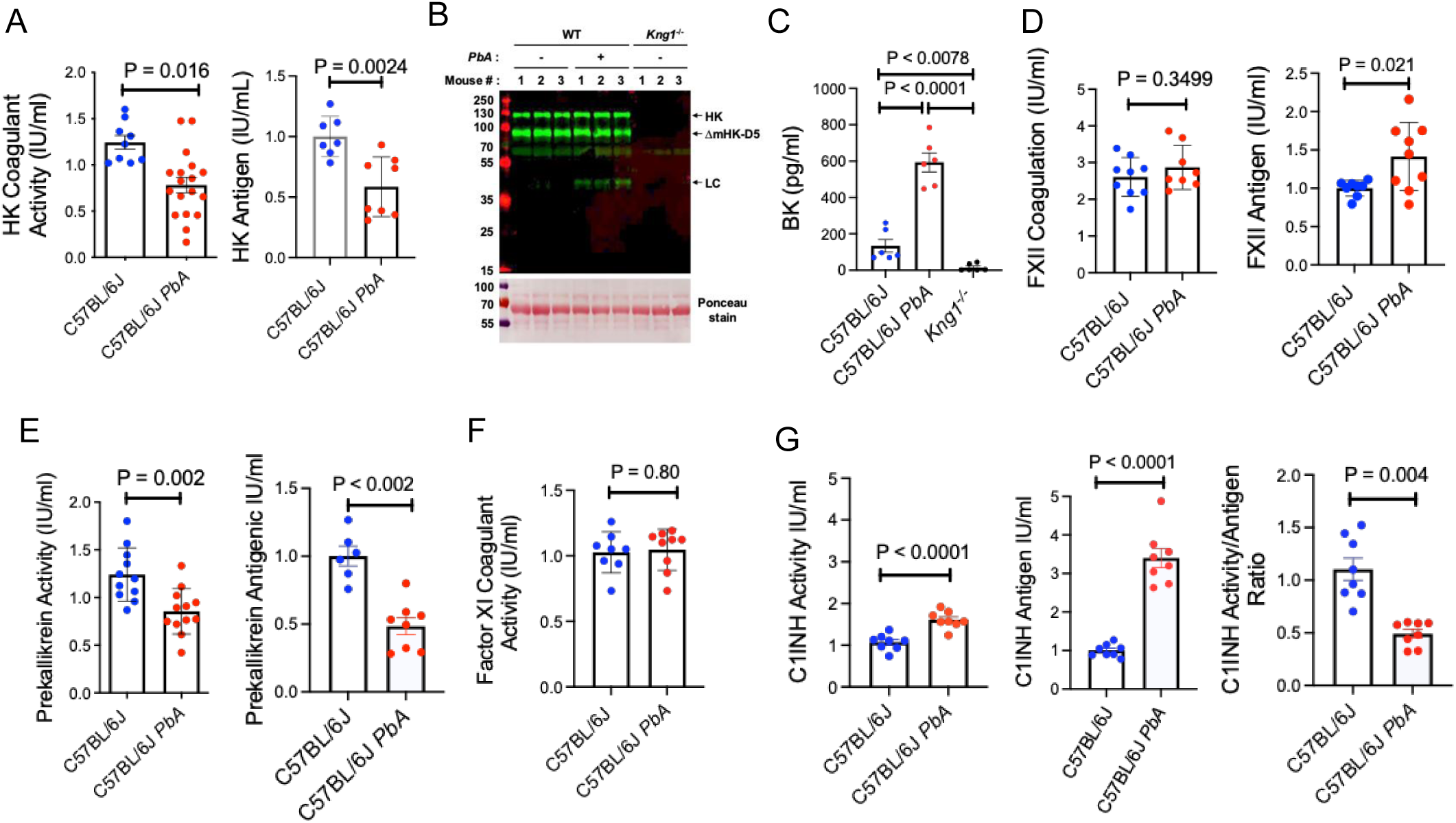
Characterization of murine plasma HK, PK, FXI, FXII, and C1INH in *PbA*-induced cerebral malaria. Panel A. HK activity was measured by coagulant assay in plasma collected into 3.2 gm% sodium citrate from uninfected (C57BL/6J) or *P. berghei* ANKA-infected (C57BL/6J *PbA*) mice. Relative murine HK antigen amount on immunoblot was determined by densitometry of equal volumes of plasma from individual uninfected or infected mice. All values presented are the mean ± SEM. Panel B. An immunoblot of a 10% Tris-Glycine gradient SDS-PAGE using a rabbit antibody reared to a sequence of murine HK domain 6. There are three reduced samples of uninfected murine plasma (*PbA-*) and three plasma samples of *PbA*-infected (*PbA* +) C57BL/6J mice. All *PbA*-infected mice manifested significant neurologic deterioration on day 5 at the time of plasma collection. The immunoblot also has three reduced samples of plasma from total kininogen deficient mice (*Kng1^-/-^*). HK represents intact high molecular weight kininogen at ∼120 kDa; ΔmHK-D5 represents of a form of mouse HK missing domain 5 (29); LC represents the cleaved 46 kDa light chain of HK. The loading control is the Ponceau stain at the bottom of the immunoblot. Panel C. Bradykinin assay by LC/MS/MS. Three kinds of samples were measured: baseline bradykinin levels in normal C57BL/6J mice, C57BL/6J mice with *PbA* infection, and total kininogen *Kng1^-/-^*mice. Panel D. Factor XII (FXII) coagulant activity was determined in individual plasmas from uninfected and *PbA*-infected C57BL/6J mice using a FXII standard curve made from a pool of plasma from healthy uninfected mice. The relative murine FXII antigen amount on immunoblot (right side) was determined by densitometry of equal volumes of individual plasma samples from uninfected and *PbA*-infected C57BL/6J mice. Panel E. Plasma prekallikrein (PK) activity assay (left side) was performed by chromogenic assay. Relative murine PK antigen amount on immunoblot (right side) was determined by densitometry of equal volumes of individual plasma samples from uninfected and Pb-A infected C57BL/6J mice. Panel F: Factor XI (FXI) coagulant activity was determined in plasma from uninfected and *PbA*-infected C57BL/6J mice against a pool of plasma from healthy uninfected mice. Panel G. C1INH functional activity (left side), C1INH antigen by immunoblot (middle), and ratio of C1INH activity to antigen (left) were determined in plasma from uninfected and *PbA*-infected C57BL/6J mice. All values shown are (mean ± SEM). For HK, PK, FXII, FXI and C1INH assays, all values are normalized to pooled normal mouse plasma and expressed IU/ml which is the amount of that protein in 1 ml plasma. Comparison between uninfected and infected mouse samples were made by two-tailed t test, P < 0.05 was considered significant.

Prominent bands of the 46 kDa light chain of murine HK (LC) were seen in plasma from infected mice (**Fig. 3B**). No bands were observed in plasma from murine kininogen deficient mice (*Kng1^-/-^*) (**Fig. 3B**). Using an anti-murine HK D5 antibody on the same plasma samples, HK was mainly a single ∼120 kDa band in plasma from uninfected and infected mice, but plasma from infected mice also showed prominent 46 kDa light chain of cHK bands. Thus, mice with ECM had the same cHK phenotype observed in plasma from children with CNS-M or UM due to *Pf* infection. Plasma from wild type C57BL/6J mice had a mean BK level of 134±35 (SEM) pg/ml (**Fig. 3C**), while *PbA*-infected mice exhibited elevated BK levels of 593±52 (SEM) pg/ml (P < 0.0001 vs uninfected mice). In contrast, *Kng1^-/-^* mouse plasma contained only 15±8 (SEM) pg/ml, a value below the lower limit of the standard curve of known BK values in total kininogen deficient murine plasma.

Subsequent investigations showed that murine FXII activity was not different between uninfected and *PbA*-infected mice (**Fig. 3D).** FXII antigen levels were increased, not decreased after *PbA* infection (**Fig 3D**). In contrast, plasma PK activity and antigen levels in *PbA*-infected mice were significantly lower (P = 0.002, P = 0.002, respectively) than uninfected mice (**Fig. 3E**). Factor XI coagulant activity was similar in plasma between uninfected and *PbA*-infected mice samples (**Fig. 3F**). Lastly, plasma C1INH activity was 60% higher in *PbA*-infected mice compared with uninfected animals (**Fig. 3G-left**). Notably, infected mice had C1INH antigen levels that were 3-fold higher than uninfected mice and 2-fold higher than their activity values (**Fig. 3G-middle).** Infected mice exhibited a C1INH activity to antigen ratio of 0.5 (**Fig 3G-right**). These data indicated that *PbA*-infected mice had circulating C1INH that was only 50% functional relative to uninfected mice.

*Role of HK and BK receptors in ECM.* C57BL/6J mice with deletions in each of the genes encoding the proteins of the KKS and factor XII were utilized to determine if any had protection from *PbA*-induced neurologic deterioration measured by the SHIRPA score or blood brain barrier integrity assessed by brain extravasation of Evans Blue dye, and increased overall survival. Total kininogen deficient (*Kng1^-/-^*) mice had a progressive increase in the level of parasitemia (∼8% at day 5 post-infection) like that of wild-type mice (**Fig. 4A Top**). Neurologic assessment at day 5 by SHIRPA score indicated that *Kng1*^-/-^ mice were protected against neurologic deterioration relative to wild type mice (P = 0.0063) and showed a significant reduction (P = 0.0006) in Evans Blue dye uptake into the brain. In a representative picture of a brain of a *PbA*-infected *Kng1^-/-^*mouse, the degree of Evans blue dye extravasation was markedly lower than in a *PbA*-infected wild type mouse. Furthermore, *Kng1^-/-^* mice exhibited increased survival with a mean of 7 days. One animal survived for 10 days whereas all infected C57BL/6J wild type mice died by day 6 (**Fig. 4A**, **Top**). These findings indicated that eliminating the source of BK alone, without treatment with anti-parasite drugs, delayed disease onset and increased survival, even in the absence of anti-plasmodial effects.

**Figure 4.**
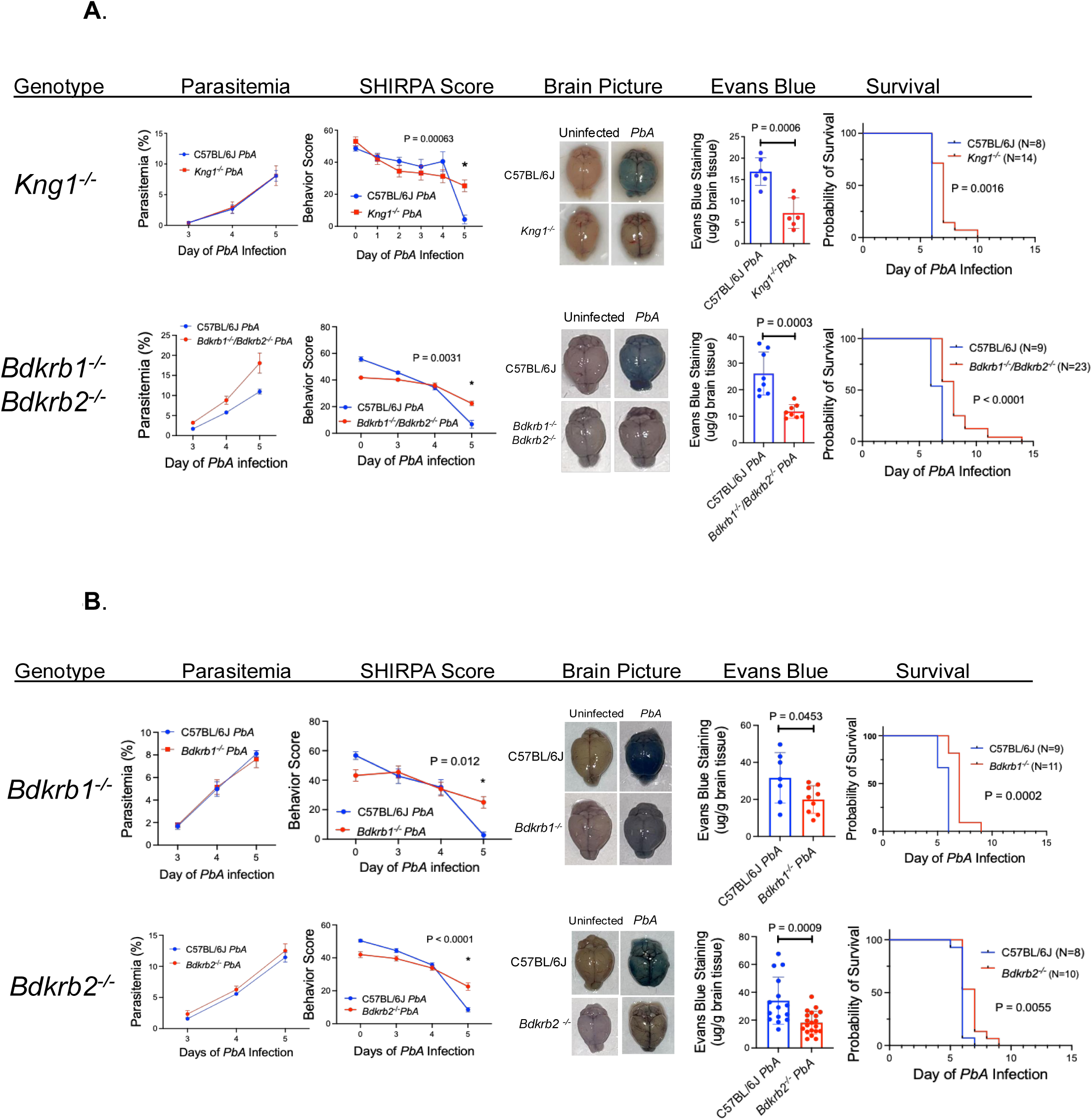
Characterization of *P. berghei* ANKA (*PbA*) infection in HK and bradykinin receptor deleted mice. The top panel characterizes kininogen (*Kng1^-/-^*) and combined bradykinin B1 and B2 receptor (*Bdkrb1^-/-^/Bdkrb2^-/-^*) null mice. **Panel A** (Top): *Kng1^-/-^* mice. Parasitemia: In *Kng1^-/^*^-^ mice (top panel, from left to right), the percent parasitemia of infected *Kng1^-/-^* mouse samples versus infected wild type C57BL/6J mice. SHIRPA Score: The parameter is a detailed motor and behavioral characterization of the mice from day 3 to day 5 before euthanasia (See Methods) (28). A score is plotted for each day 3 to 5 for infected *Kng1^-/-^*or C57BL/6J mice. Brain Picture: The brain pictures are representative photographs of wild type versus *Kng1^-/-^* mice brains after Evans Blue injection without (Uninfected) or with *P. berghei* ANKA infection (*PbA*). Evans Blue: The Evans Blue panel is a quantification of the concentration of Evans Blue dye taken up into the brain in infected wild type C57BL/6J *versus Kng1^-/-^*mice. Note: all wild type mice controls were run simultaneously with the indicated gene deleted mice. Survival: A Kaplan-Meier plot of *PbA*-infected C57BL/6J and *Kng1^-/-^* mice. The number of mice used for each group in the survival study is indicated in parenthesis on the figure’s legend. **Panel A** (Bottom): combined *Bdkrb1^-/-^/Bdkrb2^-/-^*mice were infected with *PbA* along with infection of wild type C57BL/6J controls. All lower panels are identical to the top panels describing *Kng1^-/-^*mice. Each graph is the combined data are mean ± SEM of 2 or 3 independent experiments with 5 or more mice in each experimental condition. Characterization of mice infected with *PbA* in bradykinin 1 receptor null mice (*Bdkrb1^-/-^*) and bradykinin 2 receptor null mice (*Bdkrb2^-/-^*). **Panel B** (Top): *Bdkrb1^-/-^* mice. Parasitemia: The degree of parasitemia was identical in C57BL/6J mice infected with *P. berghei* ANKA (*PbA*) (C57BL/6J) and *Bdkrb1^-/-^*. SHIRPA Score: The neurologic/behavior integrity of the mice on days 3 to 5 of *PbA* infection were determined. Brain photograph: The degree of Evans Blue uptake in the brain of *Bdkrb1^-/-^* mice versus C57BL/6J. These data were the mean ± SEM of 2-3 experiments with 5-6 mice in each. Survival: Kaplan-Meier plot of *PbA*-infected C57BL/6J and *Bdkrb1^-/-^* mice. Panel B (Bottom): *Bdkrb2^-/-^* mice. Parasitemia: The degree of parasitemia was identical in C57BL/6J mice infected with *P. berghei* ANKA (*PbA*) (C57BL/6J *PbA*) and *Bdkrb2^-/-^*. SHIRPA Score: After 5 days of *PbA* infection of C57BL/6J *PbA* versus *Bdkrb2^-/-^ PbA*). Brain photograph: The degree of Evans Blue dye uptake in the brain of *Bdkrb2^-/-^* mice compared to C57BL/6J mice. These data were the mean ± SEM of 3-4 experiments with 5-6 mice in each. Survival Kaplan-Meier plot of *PbA*-infected C57BL/6J and *Bdkrb2^-/-^* mice. Comparisons between groups on day 5 of the SHIRPA scoring and Evans blue uptake were performed by two-tailed t test, P < 0.05 was considered significant. Differences in the Kaplan-Meier plots survival curves were determined by Mantel-Cox and Gehan-Breslow-Wilcoxon tests performed independently.

Combined BK B1 and B2 receptor null mice (*Bdkrb1^-/-^/ Bdkrb2^-/-^)* (**Fig. 4A Bottom)** had a higher level of parasitemia at day 5 post-infection higher (17%) compared to control wild-type mice (10%).

However, the SHIRPA scores and Evans Blue dye uptake showed a significant reduction in neurologic deterioration (P = 0.0031) and brain swelling (P = 0.0003), respectively, relative to that in the *PbA*-infected wild type mice. In survival studies, while all *PbA*-infected C57BL/6J wild type mice all died by day 7, the double BK receptor knockout mice showed an increased mean survival of 8 days, with one animal surviving to day 14 (**Fig. 4A, Bottom**).

Next we examined murine deletion mutants from each of the two BK receptors. Parasitemia levels of control wild-type and BK B1 receptor knockout (*Bdkrb1^-/-^*) mice were identical at day 5 (**Fig. 4B Top**). However, SHIRPA scores showed a reduction in neurologic deterioration (P < 0.012) and brain Evans Blue dye uptake (P = 0.045) in knockout mice. *Bdkrb1^-/-^* mice had increased mean survival 7 days vs 6 days for wild-type mice (**Fig. 4B Top**). *Bdkrb2^-/-^*mice had a greater degree of significance from neurologic deterioration (P < 0.0001) and reduction (P = 0.0009) in Evans Blue dye uptake (**Fig. 4B Bottom**). *Bdkrb2^-/-^* mice overall survival also was 7 days (**Fig. 4B Bottom**).

*Plasma kallikrein is causal in ECM.* Since plasma kallikrein is the major enzyme that liberate BK from HK in human plasma, prekallikrein null mice *(Klkb1*^-/-^) were investigated. Parasitemia levels in *Klkb1^-/-^*mice were like that of wild-type mice. Notably, *Klkb1^-/-^* mice had a highly significant reduction in neurologic deterioration (P = 0.0003) and brain Evans Blue dye uptake (P < 0.0001) (**Fig. 5 Top**).

**Figure 5.**
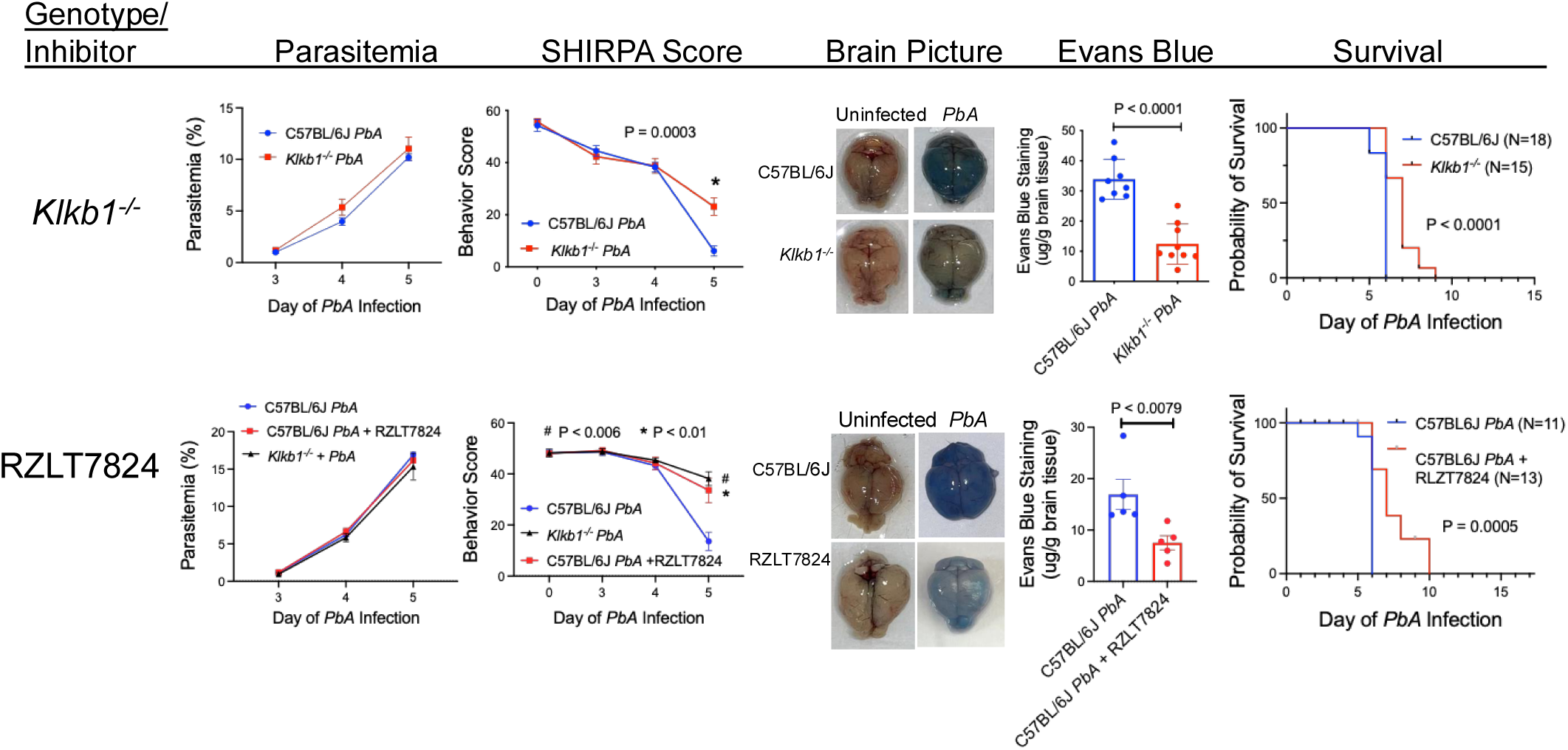
Characterization of *P. berghei* ANKA (*PbA*) infection in prekallikrein deficient mice (*Klkb1^-/-^*), and *PbA*-infected C57BL/6J mice treated with the plasma kallikrein inhibitor RZLT7824. *Klkb1^-/^*^-^ mice (Top). Parasitemia: In *Klkb1^-/^*^-^ mice (top panel, from left to right), the percent parasitemia of infected *Klkb1^-/-^* mouse samples versus infected wild type C57BL/6J mice. SHIRPA Score: This plot details the motor and behavioral characterization of the mice from day 3 to day 5 before euthanasia (See Methods). A score is plotted for each day 3 to 5 for *PbA*-infected-*Klkb1^-/-^* or-C57BL/6J mice. Brain Picture: The brain pictures are representative photographs of wild type versus *Klkb1^-/-^* mice brains after Evans Blue injection without (Uninfected) or with *P. berghei* ANKA infection (*PbA*). Evans Blue: The Evans Blue study quantifies the amount dye taken up into the brain in infected wild type C57BL/6J *versus Klkb1^-/-^* mice. Survival: A Kaplan-Meier plot of *PbA*-infected C57BL/6J versus *Klkb1^-/-^* mice. Mice treated with RZLT7824 (Bottom). *PbA*-infected C57BL/6J mice were treated with the plasma kallikrein inhibitor RZLT7824 starting on day 3. *PbA*-infected C57BL/6J mice without and with RZLT7824 were run simultaneously. The parasitemia and SHIRPA score plots compare untreated and RZLT7824-treated infected animals. A representative brain picture is shown. The Evans Blue figue compares *PbA*-infected C57BL/6J mice without or with RZLT7824 treatment. In each graph the combined data are mean ± SEM of 2 or 3 independent experiments with 5 or more mice in each experimental condition. The statistical analysis on the panels were performed as in Figure 4.

*Klkb1^-/-^* mice also showed a mean survival of 7 days whereas all the wild-type mice died by day 6 (**Fig. 5 Top**).

As an independent approach to studies of mice with genetic deletions, we determined if a parenteral treatment with a plasma kallikrein (PKa) inhibitor protected wild-type mice from ECM pathogenesis. RZLT7824 is a small molecule inhibitor of PKa with an IC_50_ of 13 nM for isolated human PKa and > 3.0 mM for human FXIIa, FXIa, FXa, FIIa, or plasmin. It inhibited mouse plasma PKa with an IC_50_ of 5 nM. RZLT7824 was administered at 60 mg/kg IP on day 3 post-*PbA* infection. Treatment with RZLT7824 preserved neurologic and motor function (P < 0.01) and reduced Evans Blue dye accumulation in the brain (P < 0.0079) on day 5 whereas untreated infected C57BL/6J mice manifested neurologic defects. RZLT7824-treated mice also had a significantly increased (P = 0.0005) mean survival of 7 days whereas all the wild-type mice died by day 6 (**Fig. 5 Bottom**). These findings with a pharmacologic inhibitor of plasma kallikrein were like those of the plasma prekallikrein null animal.

*Potential mechanism(s) of plasma prekallikrein activation in ECM.* Initial studies determined if the parasite cysteine protease, berghepain-2 has a role in BK formation in ECM. Since HK is a substrate of the cysteine protease falcipain-2 (PF3D7 1115700) (30), the contribution of the *PbA* ortholog berghepain-2 (PBANKA_0932400) to murine ECM was examined. C57BL/6J mice were infected with wild type *PbA* (*PBANKA^+/+^*) or *PbA* that had the berghepain-2 gene genetically silenced (*PBANKA^-/-^*). The levels of parasitemia were similar (10%) in the two groups at day 5. The absence of berghepain-2 expression in blood-stage parasites did not alter neurologic deterioration and Evans Blue dye brain uptake on day 5. Thus, berghepain-2 did not contribute to ECM pathogenesis.

In the contact activation system of blood coagulation, activated factor XII (FXIIa) converts zymogen PK to the enzyme PKa. We therefore infected Factor XII null mice (*F12^-/-^*) and compared them to infected wild type mice. Parasitemia levels in wild-type and *F12^-/-^* mice were similar. However, FXII deficiency did not protect infected mice from progressive neurologic deterioration. In the Evans Blue dye assay, there was a significant reduction in brain edema (P = 0.0194) but not to the extent seen with *Kng1^-/-^*, *Bdkrb2^-/-^*, or *Klkb1^-/-^*mice. This result was explained by the reduction in a boost to PK activation by the absence of FXIIa in reciprocal contact activation. Murine FXII deficiency did confer an increase in survival in ECM. *F12^-/-^* mice had a mean survival of 7 days, 24 h longer than wild-type mice.

*Prolylcarboxypeptidase is the PK activator in ECM*. Since FXII appeared to have little influence in ECM, we sought another PK activator. Plasma PK is activated on endothelial cell membranes by the serine protease prolylcarboxypeptidase (PRCP) (31). *Prcp*^gt/gt^ mice are gene trap hypomorphs with 3% PRCP mRNA (32,33). *Prcp^gt/gt^* mice had a significant reduction in neurologic deterioration (P < 0.0067) and Evans Blue dye uptake (P < 0.0007) compared with wild type controls (**Fig. 6 Top**). The mean survival of these animals after *PbA*-infection was 8.5 days (**Fig. 6 Top**). To confirm a role for PRCP in ECM pathogenesis, wild type C57BL/6J mice were treated with an inhibitor of PRCP, named PrCP inhibitor, beginning at day 3 post *PbA* infection (34). Treated mice showed a significant reduction in neurologic deterioration (P < 0.0095) and brain Evans Blue dye uptake (P = 0.039) (**Fig. 6 Bottom**). These combined findings suggest that vessel wall PRCP, a PK activator, contributed to ECM pathogenesis along with HK, BK receptors and PK independent of FXII.

**Figure 6.**
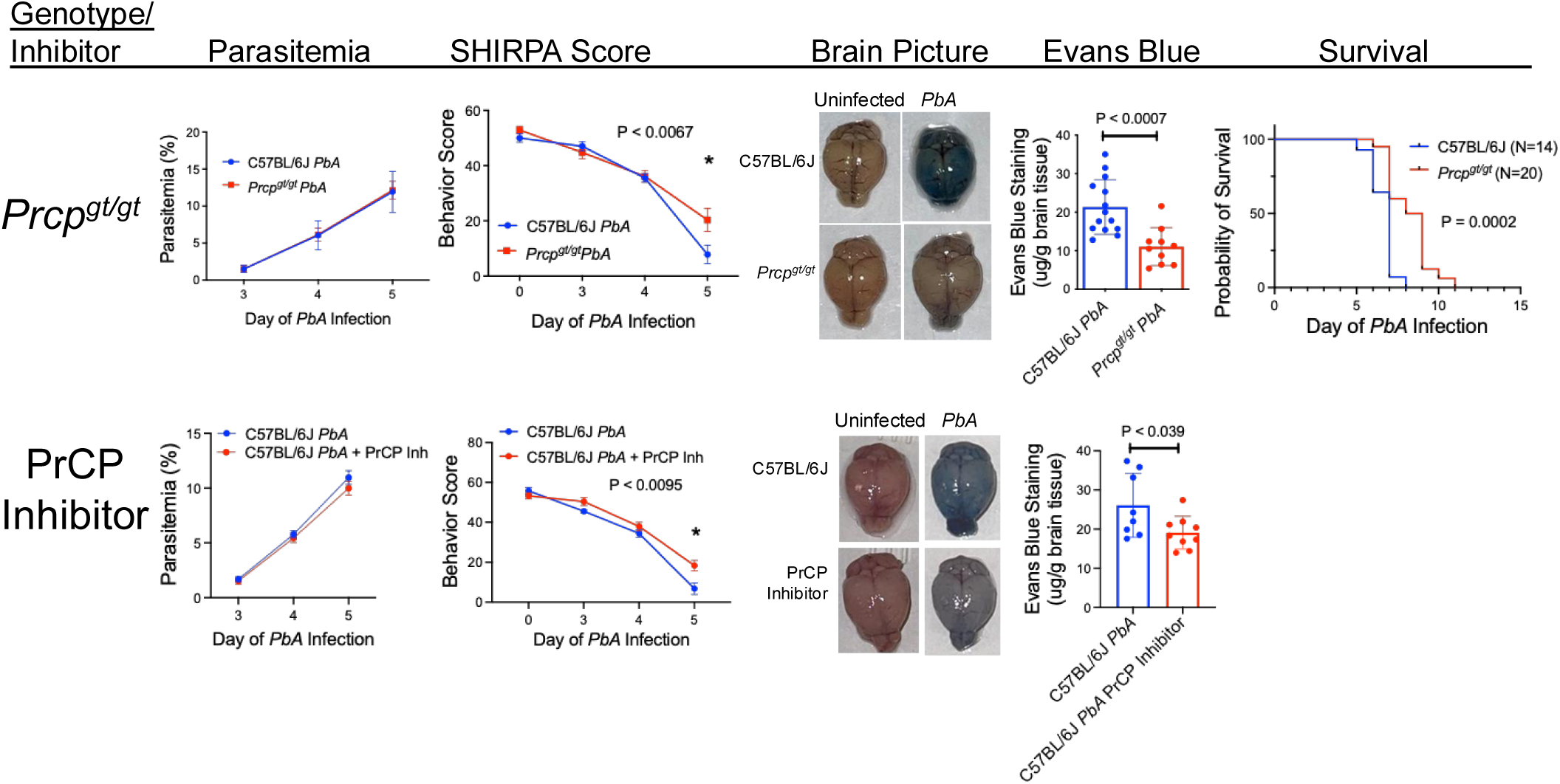
Characterization of *P. berghei* ANKA (*PbA*) infection in proplylcarboxypeptidase gene trap mice (*Prcp^gt/gt^*), and *PbA*-infected C57BL/6J given a prolylcarboxypeptidase inhibitor, PrCP Inhibitor. *Prcp^gt/gt^* mice (Top). Parasitemia: In *Prcp^gt/gt^*mice (top panel, from left to right), the percent parasitemia of infected *Prcp^gt/gt^*mouse samples versus infected wild type C57BL/6J mice. SHIRPA Score: This plot details the motor and behavioral characterization of the mice from day 3 to day 5 before euthanasia (See Methods). A score is plotted for each day 3 to 5 for *PbA*-infected - *Prcp^gt/gt^* or-C57BL/6J mice. Brain Picture: The brain pictures are representative photographs of wild type versus *Prcp^gt/gt^* mice brains after Evans Blue injection without (Uninfected) or with *P. berghei* ANKA infection (*PbA*). Evans Blue: The Evans Blue panel quantifies the dye taken up into the brain in infected wild type C57BL/6J *versus Prcp^gt/gt^* mice. Survival: A Kaplan-Meier plot of *PbA*-infected C57BL/6J versus *Prcp^gt/gt^*mice. Mice treated with PrCP Inhibitor (Bottom). *PbA*-infected C57BL/6J mice were treated with the PrCP Inhibitor starting on day 3. *PbA*-infected C57BL/6J mice without and with PrCP inhibitor were run simultaneously. The parasitemia and SHIRPA score plots compare the 2 types of infected animals. A representative brain picture is shown. The Evans Blue graph compares *PbA*-infected C57BL/6J mice without or with PrCP Inhibitor treatment. In each graph the combined data are mean ± SEM of 2 or 3 independent experiments with 5 or more mice in each experimental condition. The statistical analysis on the panels were performed as in Figure 4.

Since PRCP had not been shown to activate plasma kallikrein/kinin *in vivo*, we created a conditional *Prcp*^fl/fl^ mouse and crossed it with *Slco1c1-CreER^T2^* mice, which directs the *Cre* recombinase to the thyroxine transporter in brain microvascular endothelium (35). This *Cre* mouse has been validated by several laboratories (35–37). The loxP sites to create the *Prcp* floxed gene was placed around exon 4 (**Fig. 7A).** Homozygous *Prcp^fl/fl^* mice were crossed with hemizygous *Slco1c1-CreER^T2^*mice. Mice (named *Prcp^fl/fl^ Cre*) that had the two separate genotyping features of being homozygously floxed (*Prcp^fl/fl^*) and containing the *Cre* recombinase gene (see **Fig. 7B)** were selected for treatment with tamoxifen. Immunofluorescence studies showed that after tamoxifen treatment, *Prcp^fl/fl^ Cre* mice showed an absence of brain microvascular endothelial cell PRCP when compared *with Prcp^fl/fl^* mice, the experimental control (**Fig. 7C**). As shown in **Fig 7C**, the brain cortical tissue from *Prcp^fl/fl^* Cre mice showed strong CD31 vessel staining, but, in contrast, the green PRCP antigen was absent from the vasculature (outlined in white dotted area in **Fig. 7C Inset**). Furthermore, the adjacent line graph showed no overlap between the PRCP and CD31 antigen signals (**Fig.7C**, bottom right). To confirm the specificity of this model, renal tissue from the same mouse was examined. Both *Prcp^fl/fl^*and *Prcp^fl/fl^ Cre* mice exhibited overlapping CD31 and PRCP antigen signals in renal peritubular capillaries after tamoxifen treatment. Collectively, these data confirmed that tamoxifen-induced PRCP deficiency in *Prcp^fl/fl^* Cre mice was specific to the brain endothelium.

**Figure 7.**
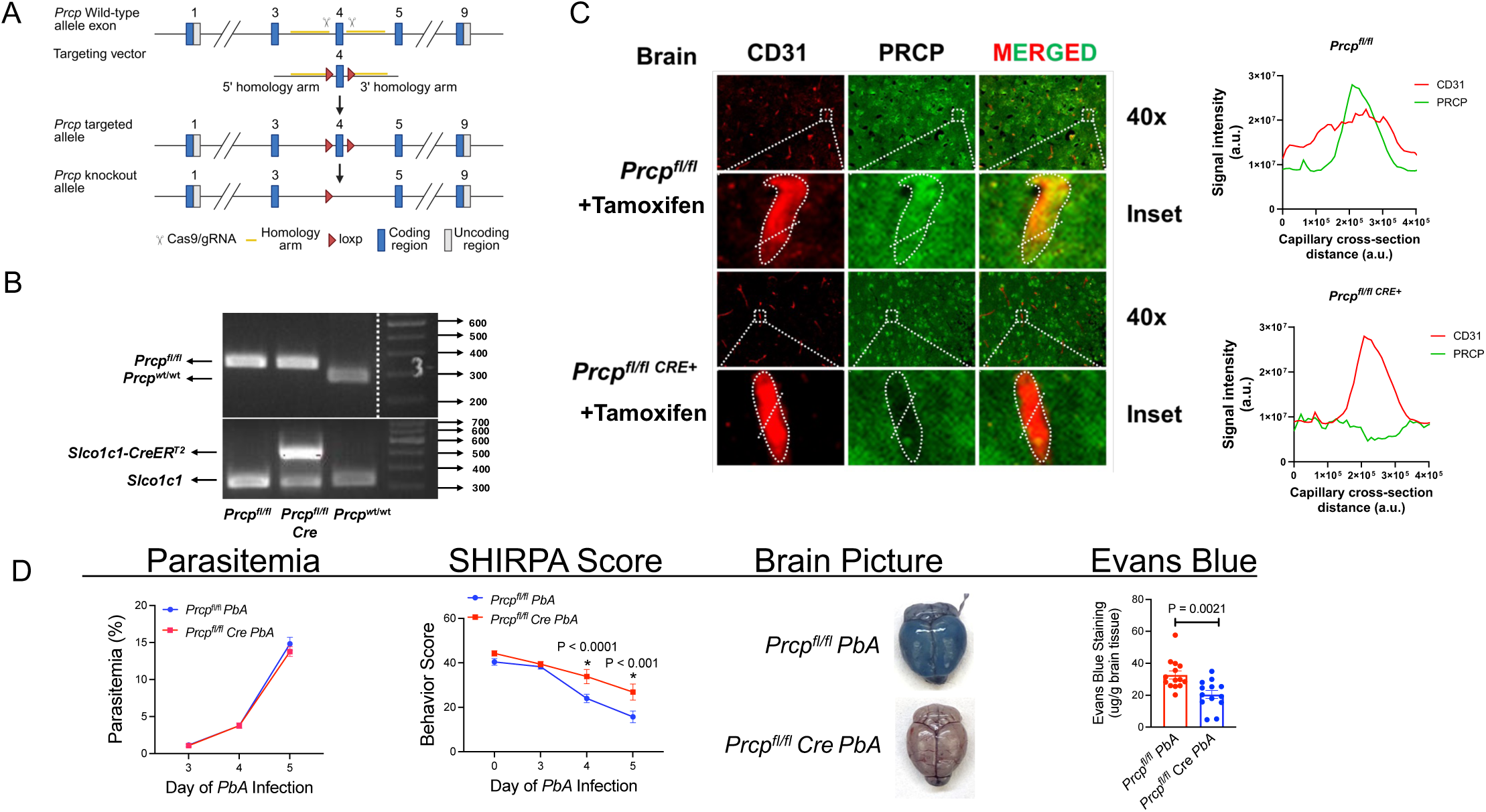
Studies with *Prcp^fl/fl^ with Slco1c1-CreER^T2^* mice. Investigations determined if a conditional knockout of brain endothelium *Prcp* would influence the *PbA* infection in C57BL/6J mice. Panel A. Targeting design for creation of *Prcp^fl/fl^* mice. The loxP sites were inserted around exon 4. Panel B. Genotyping characterization of the conditional knockout of brain endothelial cell *Prcp*. Homozygous *Prcp^fl/fl^* mice were mated with hemizygous *Slco1c1-CreER^T2^*mice to create the conditional KO mice. Panel C. Immunofluorescence of mouse brain microvascular tissue with CD31 and PRCP at 40 x and enlarged brain microvessels in the Inset. The graphs to the right of the immunofluorescence represent the cross expression of CD31 and PRCP antigen in the vessel wall. In *Prcp^fl/fl^* mice, CD31 and PRCP antigen colocalize in the immunofluorescence picture and corresponding graph to the right. In the *Prcp^fl/fl^ Cre* mice, there is no expression of PRCP antigen in the vessel (middle panel) on the immunofluorescence designated by CD31 (left panel) and in the corresponding graph to the right (there is on a flat green line for PRCP where there is a peak for CD31 expression). Panel D. *PbA* infection in *Prcp^fl/fl^*and *Prcp^fl/fl^ Cre* mice are compared. Their parasitemia, SHIRPA score, brain picture, and Evans Blue studies are shown. Significant differences on the SHIRPA score on a day and Evans Blue uptake were determined by two-tailed t test. A P < 0.05 was considered significant.

How the *PbA*-infected conditional brain-restricted microvessel *Prcp* knockout mice responded to infection is shown in **Fig. 7D**. After tamoxifen treatment, the *Prcp^fl/fl^* (the control mice for this experiment) and *Prcp^fl/fl^ Cre* mice have similar levels of parasitemia. However, there was significant protection from neurologic deterioration at day 4 (P < 0.0001) and day 5 (P < 0.001) in the *Prcp^fl/fl^ Cre* mice compared to the *Prcp^fl/fl^* mice. Further, there was a reduction of brain Evans Blue uptake (P < 0.0029) in the *Prcp^fl/fl^ Cre* mice (**Fig.7D**). These combined data indicate that brain vessel wall PRCP had a significant role in the pathogenesis of ECM.

*The use of plasma kallikrein inhibition as an adjuvant to artesunate in the treatment of experimental cerebral malaria.* Using *PbA*-infected wild type mice, we mimicked a situation that is commonly seen in hospitals in rural Africa when children present with coma and seizures with evidence of brain swelling and malaria retinopathy. We performed these studies with the PKa inhibitor RZLT7824 because 1) PK deficiency is a clinically benign condition, 2) *Klkb1^-/-^* mice are protected from thrombosis (39), and 3) various PKa inhibitors are FDA-approved to treat HAE without long-term consequences. Alternatively, PRCP is not an ideal target for these studies because its depletion produces hypertension, cardiac disease and thrombosis (33,40). The treatment protocol in **Fig. 8A** shows that on day 5 post-infection, when the animals showed objective signs of neurologic deterioration, one group was treated with the anti-parasitic drug artesunate (32 mg/kg IP) once and a second group was treated with artesunate plus RZLT7824 (80 mg/kg IP) twice, once at the same time as artesunate and 12 h later. Mice that survived 24 h from the start of treatment were evaluated. The two groups had similar parasitemia levels with a peak on day 5 before artesunate administration and ∼50% decreased parasitemia 24 h after treatment (**Fig. 8B**). When the animals’ behavior was characterized by the SHIRPA Score (**Fig. 8C**), the artesunate + RZLT7824 group had a significantly improved (P < 0.0002) SHIRPA score on day 6, indicating neurologic and motor improvement. In contrast, the artesunate alone group exhibited continued neurologic deterioration on day 6 (**Fig 8C**). Further, the combined artesunate and RZLT7824 treated animals had a reduction in brain Evans Blue dye uptake (P < 0.008) and brain mass (P < 0.0014), (**Figs. 8D & 8E**). **Fig. 8F** showed overall survival. Mice treated with artesunate alone had only 38% survival at 24 h post-treatment compared to 65% survival of the mice treated with combined artesunate and RZLT7824. Adding the PKa inhibitor to treatment with anti-parasite agent increased survival by 71%.

**Figure 8.**
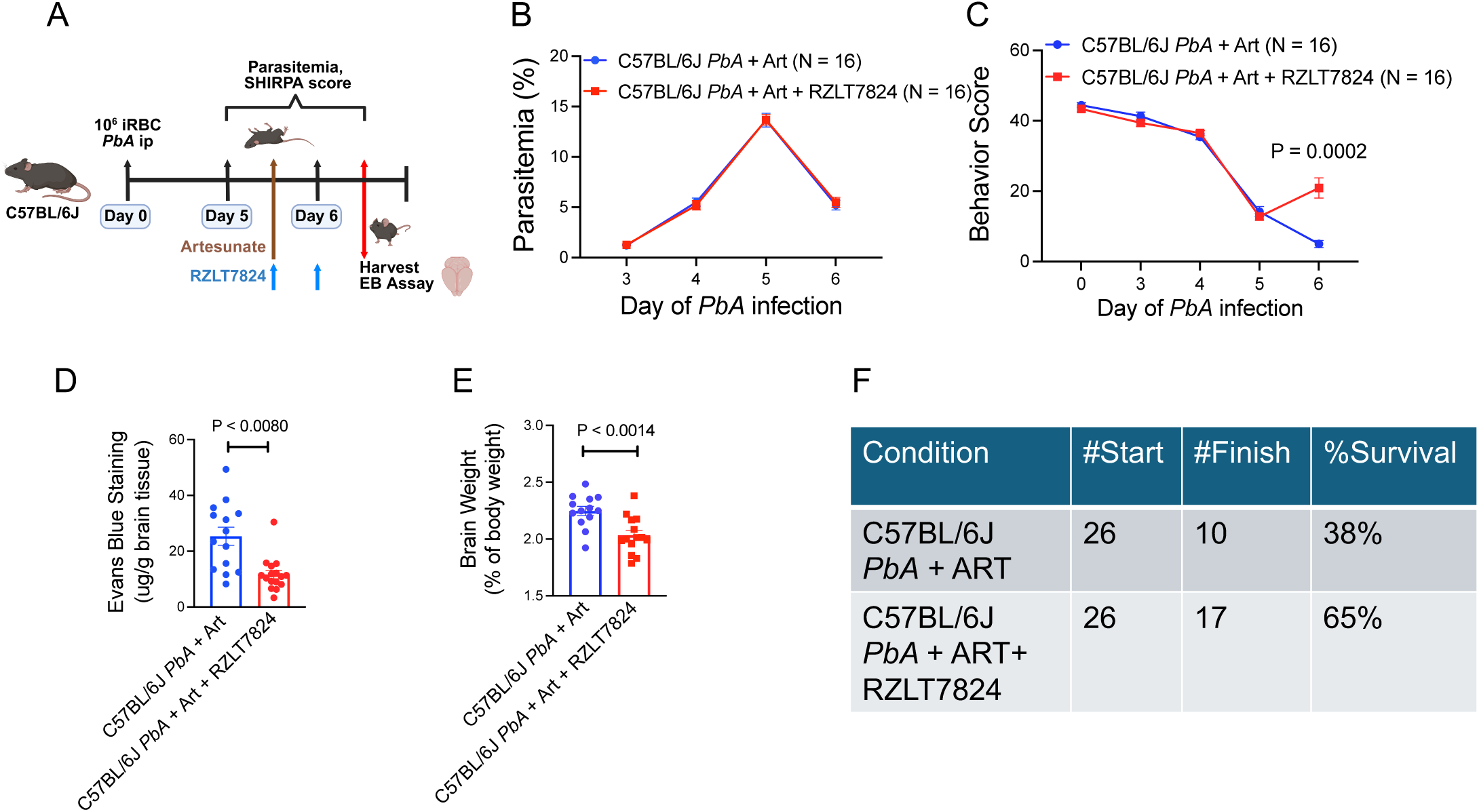
Characterization of RZLT7824 as an adjunct therapy to artesunate treatment in experimental cerebral malaria. Panel A. Schema of treatment. On day 5 when C57BL/6J mice begin to show neurologic deterioration from *PbA* infection some mice are given artesunate at 32 mg/kg IP once verses another group that gets artesunate and RZLT7824 at 80 mg/kg IP twice 12 h apart. At 24 h, the mice are assessed. Panel B. Parasitemia. Panel C. SHIRPA (behavior) Score. Panel D. Brain Evans Blue. Panel E. Brain weight. Panel F. Survival. Differences between groups were determined by two-tailed t test, P < 0.05 was considered significant.

## Discussion

Magnetic resonance imaging studies have shown that a major feature of pediatric CM is progressive brain swelling due to breakdown of the blood-brain barrier and vasogenic edema that can ultimately lead to brainstem herniation and death (7). We found that children with CNS-M and mice with ECM have cHK circulating in plasma with the presence of the 56 kDa and 46 kDa light chains of HK (**Figs. 1C and 3B**). This pattern of cleavage is an established marker of prior BK liberation, a biologic peptide that induces vasogenic edema (19–24,34). The observed HK cleavage, consistent with proteolysis by PKa or activated FXIIa indicates that the plasma contact and/or kallikrein/kinin system activation participates in the pathogenesis of CM (19–22).

The presence of circulating cHK is uncommon in humans and mice. In humans, cHK in plasma is characteristic of acute attacks of HAE due to C1INH deficiency or dysfunction (20–22). It is also seen in the plasma and brains of patients with Alzheimer’s disease and cancer (42,43). With respect to human CNS-M, only 40% of CNS-M patients and 18% of UM patients described here had detectable plasma cHK. Thus, plasma cHK by itself is not a sensitive markers of pediatric CM. *Serping1^-/-^*mice which lack C1INH protein expression, constitutively generate cHK (41). In this study, all mice with ECM had plasma cHK detectable on day 5 post *PbA* infection coincidental with the onset of objective neurologic deficits. However, the apparent differences between humans with CM and mice with ECM are not necessarily due to an inconsistency in malaria pathogenesis. In ECM, plasma is collected when all the mice have advanced neurologic signs and a reduced SHIRPA score. In real world clinical settings in rural Africa, parents bring their children to healthcare facilities at variable times after the onset of febrile illness – some at the first sign of sickness and others with more advanced disease characterized by seizures and coma. CM occurs in the course of a growing *Pf* biomass when the conserving mechanisms to prevent BK accumulation are saturated (45).

The vasogenic peptide implicated in this study as a primary mediator of ECM is elevated levels of BK. Further, genetic deletion of murine HK, the precursor of BK, provided strong protection against ECM pathogenesis. *PbA*-infected BK receptor-deleted (combined *Bdkrb1^-/-^ /Bdkrb2^-/-^*, *Bdkrb2^-/-^*, or *Bdkrb1^-/-^*) mice also had reduced ECM severity with protection from neurologic deterioration and brain Evans Blue dye extravasation, corroborating the assessment that the biologically active peptide BK contributes to vasogenic edema. It is important to appreciate that the effect of these knockout mice on ECM was not directed at the parasite and only influenced cerebral edema formation. Experimental studies with cultured human brain vascular endothelial cells have shown that both the B2 and, to a lesser extent, B1 BK receptor, contribute to loss of barrier function, a critical step in vasogenic edema formation (15). In addition to stimulation of its own receptors, local BK formation may be proximal to barrier function changes via the VEGF-VEGFR2 and angiopoietin 1-TIE2 systems. BK stimulates VEGF production which is elevated in CM and the BK B2 receptor transactivates the VEGFR2 (KDR/Flk-1) (46–48). BK also mediates its effect on barrier function through interacting pathways with angiopoietin 1-TIE2. Angiopoietin-2 null mice block BK-stimulated vascular leakage (49). Moreover, over expression of angiopoietin-1 blocks BK-induced vascular leakage and blockage of angiopoietin-1 promotes it (50). Additional studies are needed to examine the connections between the BK receptors and VEGF/VEGFR2 and the TIE2-angiopoietin systems.

In mice, we identified a novel mechanism for activation of the kallikrein/kinin system in ECM. Since the *Pf* protease falcipain-2 cleaves HK to produce cHK and BK (30), we were surprised to observe that *PbA*-infection with trophozoites that did not express the falcipain-2 ortholog berghepain-2 were not protected from CM. Further, we did not anticipate *F12^-/-^*mice would have the least protection from ECM compared to all other murine deletion models examined. We speculate that the minor impact of FXII deletion on Evans Blue dye extravasation observed is the result of an intact PK activation system that did not have a burst in PKa formation by reciprocal activation of formed factor XIIa. Alternatively, PK deficiency was highly protective against ECM and led us to consider another PK activator independent of FXII. Prolylcarboxypeptidase (PRCP) is an endothelial cell S1 serine protease that is an activator of PK to PKa independent of FXII (31,51). We observed that *Prcp^gt/gt^* mice, C57BL/6J wild type mice treated with a PRCP inhibitor (34), and *Prcp* brain microvascular conditional knockout mice had reduced neurologic deterioration and Evans Blue dye extravasation. Endothelial cell PRCP activates PK to PKa independent of FXIIa. The activated PKa then cleaves HK to liberate BK, which binds both the B2R and the B1R and produces vasogenic edema (**Figure 9A**) (34). This pathway is initiated at the vessel wall and may be unique to murine ECM since microvascular vessel occlusion is the observed mechanism (52)

**Figure 9.**
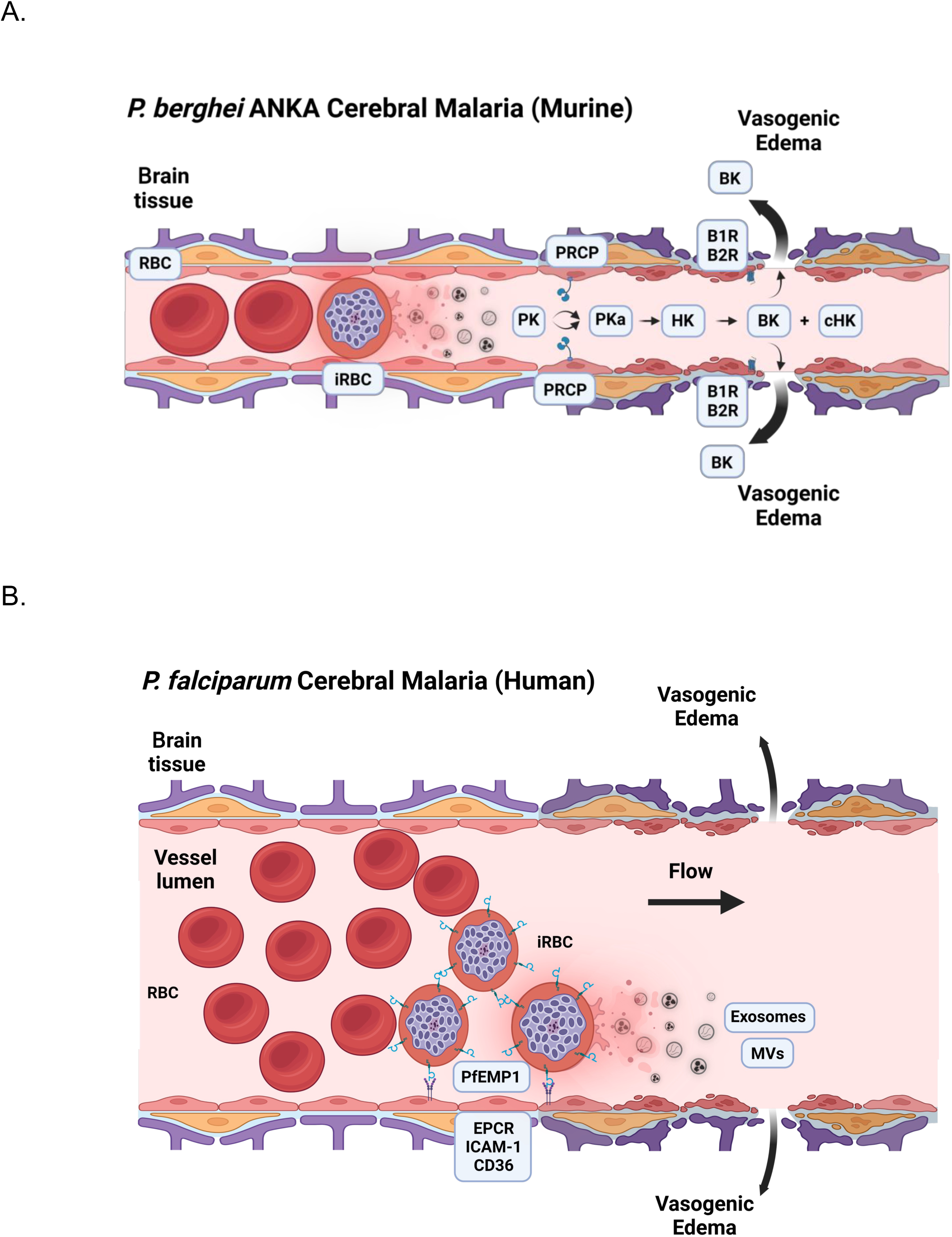
Panel. **A.** *P. berghei* ANKA (*PbA*) model of murine cerebral malaria. In *PbA*, single cell brain microcapillaries became occluded with infected red blood cells (iRBC). The occlusion of the vessel and or the effect of iRBC material plugging the vessel influences its endothelium to allow vessel wall proplylcarboxypeptidase (PRCP) to activate murine plasma prekallikrein (PK) to plasma kallikrein (PKa). PKa cleaves the HK bound to it to produce increased cHK and liberating bradykinin (BK). Formed BK binds to the bradykininB2 receptor (B2R) and the bradykinin B1 receptor (B1R). The B2R was the predominant receptor for BK binding. Activation of the B2R contributes to reduced endothelial cell barrier function and vasogenic edema. Gene deleted mice, *Kng1^-/-^*, *Klkb1^-/-^*, *Prcp^gt/gt^*, combined *Bdkrb1^-/-^* /*Bdkrb2^-/^*^-^, *Bdkrb2^-/-^*, *Bdkrb1^-/-^* and *Prcp^fl/fl^ Cre* mice significantly reduce brain edema and protected the animals from neurologic deterioration. Alternatively, *F12^-/-^* mice do not significantly protect the mice from neurologic deterioration. These data indicate that the plasma kallikrein/kinin systems contributed to the vasogenic edema in *PbA*-induced murine (experimental) cerebral malaria. **Panel B**. Human cerebral malaria pathogenesis. Unlike mice, human CM occurs in larger post-capillary venules where single iRBCs do not occlude the vessel. *Pf*-infected RBCs express on their surface a protein called PfEMP1 (Plasmodium falciparum membrane protein1) that binds the iRBCs to the vessel wall by the endothelial cell membrane protein that include endothelial cell protein C receptor (EPCR), intercellular adhesion molecule 1 (ICAM-1), and CD36. BK liberation and vasogenic edema could be initiated by adherent iRBCs themselves or others that have ruptured. Microvesicles and/or exosomes also could serve as a potential source for falcipain-2 and indirectly generate BK through surface contact activation of factor XII by autoactivation and/or stimulate the endothelial cell membrane-associated enzyme prolylcarboxypeptidase. Both enzymes lead to activation of prekallikrein and the plasma kallikrein/kinin system. Both figures were produced with BioRender.

Our ECM studies suggest that the entire plasma kallikrein/kinin system, not a single substrate (e.g., HK), enzyme (e.g., PKa, PRCP), or receptor (e.g., B2R, B1R) are host factors that contribute to its pathogenesis. Elimination or interference with the function of any kallikrein/kinin system component ameliorates ECM. In theory, targeting any of these proteins by any technique could be contributory to improve the outcome of ECM. Since CM and ECM are disorders of a swelling brain within the fixed space of the skull, treatments targeting the vasogenic edema itself along with anti-parasite therapy may produce improved outcomes in children. This notion was first demonstrated in mice by Higgins SJ, et al. in their ground-breaking investigation that showed that angiopoietin 1 infusion in mice along with artesunate treatment significantly increased ECM survival (14). Our investigation on the BK system has the same conclusion on the importance of adjunct treatment in CM to decrease brain edema along with anti-parasite therapy. Artesunate-based therapy is currently the mainstay of treatment for pediatric CM. Clinical recovery is often observed within approximately 2 days, the time it takes to eliminate the parasite. In the mouse model we show that at 24 h after adding plasma kallikrein inhibition to artesunate therapy, there is 1) improvement in neurologic behavior, 2) reduction in brain Evans Blue uptake and mass, and 3) increased survival compared with artesunate therapy alone. These data indicate that in addition to ongoing efforts to eliminate malaria by vaccines and control acute disease with anti-parasite therapy, adjunctive treatment specifically targeting cerebral vasogenic edema by several pathways may substantially increase survival by more rapidly reducing brain swelling. This approach may also have potential to improve neurocognitive outcomes in surviving children.

It is well documented that human CM due to *Pf* infection and ECM due to *PbA* infection are different in several ways. *Pf*-infected RBCs are sequestered in medium-and large-sized brain vessels that express RBC surface proteins such as PfEMP-1 that bind to endothelial cell protein C receptor (EPCR), ICAM and CD36 on the vessel wall (**Figure 9B**). In contrast. *PbA* iRBCs occlude mouse brain microvascular vessels where the diameter of the arteriole or capillary is about the same size of the diameter of the RBC (**Figure 9A**) (52). Normal flexible RBCs pass through vessels readily, but infected RBCs have reduced flexibility and occlude the vessel leading to distal thrombus (**Figure 9A**). Although these features distinguish human CM and mouse ECM from each other, the final common pathway for vasogenic edema in both may be due to BK formation and similar pathophysiology in the brain. Notably, blood-brain barrier dysfunction with MRI evidence for vasogenic edema, brain accumulation of cytotoxic CD4+ and CD8+ T cells, and extravasation of both uninfected and iRBCs occur in both (12,13,52–56). If BK is pathogenetic for the vasogenic edema in *Pf* CM, formation of BK could arise from different mechanisms in human CM and mouse ECM. In human, BK formation in *Pf* could occur by falcipain-2 directly cleaving HK, FXII autoactivation on parasite, RBC, or endothelial cell extracellular vesicles activating PK leading to BK formation, or vessel wall PRCP activating PK independent of FXIIa (**Figure 9B**). However, regardless of the precise mechanisms of BK formation, if BK is present, it will induce vasogenic edema in humans through mechanisms like those observed in mice. Hence, the same therapeutic agents effective in ECM may also be applicable in humans.

*Limitations of the study.* The hypothesis that BK is a mediator of *Pf* CNS-M could not be confirmed in the Kenyan children because 1) plasma samples were collected in sodium heparin without protease inhibitors to prevent in vitro formation and degradation of BK. Further, plasma collected into sodium heparin obviated our ability to measure residual PK, FXII, or FXI in plasma and interfered with C1INH antigen studies. Additional studies are needed to determine if *PbA*-infected mice treated with artesunate plus RZLT7824) have preserved long-term neurologic and cognitive function relative to treatment with artesunate alone.

## Methods

### Sex as a biologic variable

Our study examined male and female children, and similar findings are reported for both sexes. Likewise, our study examined male and female mice, and similar findings are reported for both sexes.

### Materials

The chromogenic substrates, H-D-Pro-Phe-Arg-pNA·2HCl (S-2302) (Cat #S820340) for plasma kallikrein. αFXIIa, and βFXIIa, H-D-Glu-Pro-Arg-pNA.2HCL (S2366) (Cat # S821090) for factor XIa, H-D-Phe-Pip-Arg-pNA.2HCl (S2238) (Cat # S820324) for thrombin (FIIa), and H-D-Val-Leu-Lys-pNA.2HCL (S2251) (Cat # S820332) for plasmin were purchased at DiaPharma. Z-D-Arg-Gly-Arg-pNA 2HCL (Biophen-CS11) (Cat # A229014) for factor Xa was purchased from Anaira Diagnostica.

Purified single chain HK, PK, PKa, FXIIa, βFXIIa, FXIa, FXa, thrombin, and plasmin were purchased from Enzyme Research Laboratories (Cat # HT1300, Cat # HPK1302, Cat # HPKa1303, Cat # HFXIIa 1212a, Cat # HFXIIab, Cat # HFXIa 1111a, Cat # HFXa 1011, Cat # HT 1002a, and Cat # Hplasmin, respectively). The cysteine protease inhibitor E64 was obtained from Sigma-Aldrich (Cat #E3132). The fluorogenic substrate Z-Phe-Arg-AMC.HCl was purchased from Bachem (Cat #4003379). The PRCP inhibitor (named “PrCP Inhibitor”) was purchased from Sigma-Aldrich (Cat #5.04044). This compound is a brain-penetrant, dichlorobenzimodazolopyrrolidinamide compound with selective inhibition of PRCP of 1 to 2-8 nM of human or mouse PRCP in the absence or presence of albumin (57). RZLT7824, a plasma kallikrein inhibitor from Rezolute, Inc., was generously provided by Dr. Jeffrey Breit, Rezolute, Inc., Redwood City, CA. Artesunate was purchased from Sigma-Aldrich (Cat # PHR2573).

### Plasma preparation from malaria study participants

Peripheral blood samples from Kenyan children under 10 years with central nervous system malaria (CNS-M), uncomplicated malaria (UM), acute febrile non-malarial illness (AFN), or healthy community control children (Healthy) were collected into BD Heparin Blood Collection tubes at the Chulaimbo Sub-County Hospital in western Kenya (26). Please note that the term CNS-M, not cerebral malaria (CM), is used for these children in accordance with our recent study indicating that healthcare personnel did not have the capacity perform ophthalmoscopic examination to evaluate retinal pathology (26). Blantyre scores of children diagnosed with CNS-M ranged from zero to 3. Ten of 20 children examined in the current report had Blantyre scores of ≤ 2. Plasma was prepared from these samples and were frozen at-80°C. These samples remained frozen until assay at Case Western Reserve University. In control experiments, plasma from healthy malaria naïve North American adults was collected into sodium heparin (1 U/ml), EDTA (5 mM), or sodium citrate (3.2 gm%) anticoagulants and showed similar structural stability of plasma HK when incubated at 37°C at 1 and 24 h and room temperature for 1 week. No sample showed spontaneous cleavage of intact 120 kDa HK into 56 and 46 kDa bands on reduced SDS-PAGE.

Blood from the CNS-M patients (See **Table 1**) was collected on admission to the hospital (CNS-M Entry, CNS-M-En) (26). CNS-M diagnosis was based on clinical presentation with seizures/coma (Blantyre score 0-3), fever, a *Pf* positive blood smear, and no other apparent cause of neurologic dysfunction. A paired blood sample was collected at the time of hospital discharge (CNS-M Discharge, CNS-M-D) 2 to 3 days later following completion of treatment with intravenous artesunate, clinical recovery with Blantyre score of 5, a lack of fever, and a *Pf* negative blood smear (58). Blood was also collected from children with uncomplicated malaria (UM) (*Pf* positive smear, no CNS symptoms or signs, Blantyre score = 5), children with acute fever non-malaria illness (AFN), and healthy children living in the same community. Healthy children did not have fever (axillary temperature ≤37.8^0^C), had a *Pf* negative blood smear, and no history of clinical malaria in the previous 6 weeks. CNS-M patients were hospitalized for intravenous treatment with artesunate. UM and AFN children were treated as outpatients and did not have recovery blood samples obtained after oral treatment with artesunate or antibiotics (27).

### Animal studies

Wild type (WT) C57BL/6J (C57BL/6J) mice were purchased from Jackson Laboratories (Bar Harbor, ME). All knockout mice were in a C57BL/6J background. All mice (wild type and knockout) had ECM following infection with *Pb* ANKA. *Kng1*^-/-^ knockout animals were provided by Dr. Keith R. McCrae (29). *F12*^-/-^ mice have been previously described (58). *Klkb1*^-/-^ mice were developed at the Texas Genomic Institute and generously provided by Dr. Edward Feener, Beth Israel Hospital, Harvard University (59). *Prcp*^gt/gt^ mice were generated at the University of Michigan from KST302 ES cells obtained from Bay Genomics (32). *Bdkrb1^-/-^* mice were provided by Dr. Michael Bader (60). *Bdkrb2^-/-^*mice, strain name B6/129S7-*Bdkrb2*^tm1Jfh^ originally was purchased from Jackson Laboratories (Bar Harbor, ME). These animals were backcrossed 10 generations into a C57BL/6J background (61). The double bradykinin B1 and B2 receptor deficient mice (*Bdkrb1^-/-^/Bdkrb2^-/-^)* were kindly provided by Drs. Rafal Pawlinski and Erica Sparkenbaugh, from the University of North Carolina, Chapel Hill, North Carolina, USA (62). Immunoblots of *Kng1^-/-^, Klkb1^-/-^,* and *F12^-/-^* of plasma from each of these mice had no plasma protein on immunoblot *and* the *Prcp^gt/gt^* had no PRCP antigen on immunoblot of kidney. Purchased C57BL/6J wild type mice were used as simultaneous controls throughout these experiments.

### Preparation of Brain Endothelium PRCP Conditional Knockout Mice

*Prcp* conditional knockout mice (C57BL/6JSmoc-*Prcpem(flox)/Smoc*) (*Prcp^fl/fl^*) (Cat. NO.: NM-CKO-230233) were made at Shanghai Model Organisms Center, Inc., Shanghai, China (63). Exon 4 of the *Prcp* transcript was the floxed region (Figure 7A). The targeting construct using CRISPR/Cas9 gene editing was composed of FRT-splice acceptor-IRES-lacZ-polyA-loxP-neomycin cassette-polyA-FRT-loxP inserted upstream of exon 4. A second loxP site was inserted downstream of exon 4. Flp-mediated recombination removed the FRT-flanked lacZ and neo cassette leaving exon 4 floxed. Genomic DNA was isolated from the tail and the *Prcp^fl/fl^* mice were genotyped by polymerase chain reaction (PCR) by 32 cycles of 94°C for 30 sec, 58°C for 30 sec, and 72°C for 5 min. The forward (TCCGCCTGGCTAAATGTTCA) and reverse (CGGCAGAGGTTCAGA-ACAGA) genotyping primers defined a 248 bp band in wild type, a 343 bp band in homozygous floxed, and both bands in heterozygous mice with the floxed gene (**Figure 7B**).

To prepare the conditional knockout, *Slco1c1-CreER^T2^*mice were obtained from Professor Markus Schwaninger at the University of Lubeck, Lubeck, Germany. These mice take advantage of the selective expression of the thyroxine transporter, *Slco 1c1*, in brain endothelial cells (35). Briefly, a bacterial artificial chromosome (BAC) was chosen to contain the mouse *Slco1c1* locus and by homologous recombination the Cre recombinase (iCre) and a mutated human estrogen receptor (*ER^T2^*) were electroporated into bacteria. The cloned genomic fragment with the *iCreER^T2^* were inserted at the ATG site of the *Slco1c1* gene (35). Transgenic offspring were created by microinjection into B6D2F1 hybrid mouse pronuclei (35). Genomic DNA was isolated from the tail and the *Prcp* mice were genotyped by polymerase chain reaction (PCR) by 30 cycles of 94°C for 30 sec, 60°C for 30 sec, and 72°C for 45 sec. The forward (CTA GGC CAC AGA ATT GAA AGA TCT) and reverse (GTA GGT GGA AAT TCT AGC ATC ATC C) genotyping primers defined a Cre-324 bp band in wild type. A 521 bp band produced by the forward (GCTATTCAT-GTCTTGGAAGCC) and reverse (CAGGTTCTTCCTGA-CTTCATC) genotyping primers defines the presence of the *Cre+*.

Demonstrating both bands defines hemizygous transgenic mice containing the floxed gene (**Figure 7B**). (*Prcp^fl/fl^ Cre*+). Brain endothelial cell deletion of mouse PRCP was performed by mating homozygous *Prcp^fl/fl^*mice with transgenic hemizygous *Slco1c1-CreER^T2^* mice. The *Cre* recombinase in the homozygous *Prcp^fl/fl^ ^Slco1c1-CreERT2^* (named *Prcp^fl/fl^ Cre*) mice was induced by 50 mg/kg tamoxifen for 5 days.

Additional validation that brain endothelial cell PRCP was removed was performed by immunofluorescence. Briefly, mouse brain and kidney sections were incubated with antibodies to CD31 and mouse PRCP (See Supplemental Table S1 for complete description of the primary and secondary antibodies used in these studies). The goat anti-mouse polyclonal antibody to mouse PRCP (anti-TDN20) was reared with a peptide from the mouse PRCP amino acid sequence TNDFRKSGPYCSESIRKSWN at Q.C.B. Custom Antibody Service (33). Quantification of CD31 and PRCP expression in the brain sections and kidney were performed with assistance of FIJI software (64). Immunofluorescence micrographs channels were separated. and regions of interest (ROIs) were selected using the straight-line tool. In the “Analyze” toolbox, Plot Profile was selected. Values of distance and signal intensity were plotted in XY axis graphs using GraphPad Prism 10.6.1. The signal from the same ROI in the green and red channels were obtained and plotted as indicated. Regions where both signal intensities increase were considered as colocalization. Region where the signal intensity is flat indicates no antigen present.

### Parasites

The wild-type rodent malaria parasite, *Plasmodium berghei* ANKA, MRA-868 7 was obtained through BEI Resources. Transgenic *P. berghei* ANKA parasites (*PBANKA^-/-^*) [line 1619 (www.pberghei.eu; RMgm-32)] that lack the gene encoding Berghepain-2 (PBANKA_0932400) were obtained from Dr.

Chris J. Janse, Leiden Malaria Research Group, Leiden University Medical Center, Leiden, Netherlands (65). Both wild-type and transgenic parasite lines have been maintained by combination of passage in C57BL/6J mice and cryogenic storage of *PbA*-infected red blood cells (iRBCs) (66).

### Murine ECM

C57BL/6J wild-type and gene deficient mice were inoculated IP with 0.1 mL suspension of 10^6^ *PbA*-infected red blood cells (67). The progression of ECM was evaluated by measuring *PbA* parasitemia and motor and neurological behavior assessment (SHIRPA scores) on day 3, 4 and 5 after infection (28,67). The SHIRPA score is based on observations of the neurological behavior, including transfer arousal, locomotor activity, tail elevation, wire maneuvers, and contact righting reflex were observed (28,68). Each are scored from 0 to 40. A high number is normal. Scores in ECM mice are below 20. Parasitemia was measured by BD-LSR II (BD Bioscience) flow cytometry using an optimized Hoectsh-Thiazole Orange staining strategy which was confirmed by optical microscopy of Giemsa-stained thin blood smears by analyzing at least 10 random microscopic fields (68). Final values were expressed as the percent of infected red blood cells (68). When the PRCP inhibitor (PrCP Inhibitor) was used with wild-type mice, the animals were injected intraperitoneally daily with 100 μl of the agent starting on day 3 of the infection with a calculated dose based on the estimated blood volume (7% body weight) to achieve a peak concentration of 100 μM inhibitor (34). When the plasma kallikrein inhibitor RZLT7824 was used with wild-type mice, the animals were injected IP with 100 μl of the solution that was calculated to be 60-80 mg/kg mouse every 12 h also starting on day 3. In investigations comparing artesunate alone versus artesunate and RZLT7824 together began at day 5 when the animals had objective neurologic defects. Artesunate was administered at a dose of 32 mg/kg in 100 μl IP once daily and RZLT7824 was given twice. Note: animal movies 1 and 2 (control C57BL/6J and genetic deletion mice) were made on day 5 when the control animals demonstrated objective neurologic deterioration on the SHIRPA score. Animal movie 3 was made on day 6 after 24 h of treatment.

### Cerebral edema measurements

The evaluation of cerebral edema was performed using the Evans Blue extravasation assay (69). Briefly, on day 5 of *PbA* infection mice received an intravenous orbital injection of 1% Evans Blue dye solution when objective neurologic findings were observed in C57BL/6J wild-type mice. After 2 h, the mice were euthanized and their brain was harvested, weighed, and incubated in 1 ml of formamide (37°C, 24 h) to extract the dye. Absorbance of the supernatant was measured at 600 nm. The concentration of dye was determined using a standard curve. Data are expressed as micrograms of dye normalized per gram of tissue. Brain mass of wild type versus *PbA*-infected mice without or with treatment was determined by direct weighing of the excise brain from the cranial cavity.

### Murine complete blood counts

On day 5 of *PbA* infection, whole blood was collected from infected and control mice anesthetized with isoflurane. Following a midline celiotomy, blood was drawn from the inferior vena cava using a 20-gauge needle attached to a 1 mL syringe containing 0.05 mL of 3.2% sodium citrate.

Approximately 0.5 mL of blood was collected and centrifuged at 514 × *g* for 5 minutes at room temperature. The resulting plasma was either used immediately or stored in aliquots at –80°C. Complete blood counts were performed using a HEMAVET analyzer according to the manufacturer’s instructions.

### General coagulant assays

Activated partial thromboplastin time (aPTT) was performed by mixing 50 μl citrated plasma from infected and non-infected mice with 50 μl pre-warmed aPTT reagent (Helena Laboratories) in a glass tube and incubated for 5 min at 37°C. The reaction was initiated by adding 50 μl of 35.3 mM CaCl_2_.

The endpoint clotting time was determined visually by constantly tilting the glass tube in a 37°C water bath. The prothrombin time (PT) was performed similarly by the addition of 50 μl of pre-warmed PT reagent which contains CaCl_2_ (Thromboplastin Reagent, Helena) to 50 μl citrated plasma from infected and non-infected mice in a glass tube.

### Coagulation factor assays

HK, FXI and FXII plasma levels were measured in infected and non-infected mice using human HK-, FXI-and FXII-deficient plasmas as substrate in an aPTT-based coagulant assay, as described previously (70). William’s trait (total kininogen deficiency) human plasma donated to this laboratory was used as substrate for the HK assay. Factor XI-and XII-deficient human plasma was obtained from George King Inc, Overland Park, KS. A standard curve was created using a serial dilution from pooled normal mouse plasma to calculate the levels of these proteins in murine samples. The final values were expressed in IU/ml that is defined as the amount of factor activity or antigen in one ml pooled normal human or murine plasma.

### Prekallikrein chromogenic assay

The PK levels in plasma from infected and non-infected mice were measured by chromogenic assay (71,72). Fifty μl of each plasma sample was treated with 50 μl of 0.167 M HCl for 15 min at room temperature to eliminate C1 inhibitor and other SERPIN plasma kallikrein inhibitors (72). Then, 50 μl of 0.1 M Na phosphate, 0.15 M NaCl, 1 mM EDTA pH 7.6 buffer and 50 μl of 0.167 M NaOH were then added to neutralize the acid treatment. To determine the residual plasma PK activity, 50 μl of treated samples was incubated with 50 μl of plasma PK activator (260 pM of βFXIIa) for 5 min. The concentration of βFXIIa added to activate the plasma PK was previously shown to be insufficient to hydrolyze the chromogenic substrate itself. After the activation period, 50 μl of the chromogenic substrate S2302 (2 mM) was added, the reaction was stopped after 10 min of hydrolysis using 100 μl of 50% acetic acid and the OD was obtained at 405 nm. A standard curve was created using a serial dilution from pooled normal mice plasma. The final values are expressed in IU/ml. One unit is the amount present in 1 ml of pooled normal mouse plasma.

The specificity of RZLT7824 to inhibit isolated plasma kallikrein (PKa) and activated plasma prekallikrein in plasma was determined. The IC_50_ of 1 nM PKa or 1 nM FXIIa using 30 to 3000 nM RZLT7824 were determined using 0.4 mM S2302. The IC_50_ of 3 nM FXIa, FXa, thrombin, or plasmin using 30 to 3000 nM RZLT7824 was determined using 0.3 mM S2366, 0.15 mM Biophen-CS11, 0.3 mM S2238, or 0.4 mM S2251, respectively, in a Hepes buffer, pH 7.4.

### C1 inhibitor functional assay

Plasma C1INH functional levels were performed by the assay of Shapira *et al*. modified for a microplate reader (73). After mouse blood collection, 50 μl from infected and non-infected mouse plasma samples were first treated with methylamine (40 mM final concentration) for 2 h at 23°C to eliminate plasma α_2_macroglobulin (74). Ten μl of the treated samples were diluted with 50 μl in PBS buffer containing 2 mg/mL of bovine serum albumin (fraction V). The reaction was started by adding 10 μl of plasma kallikrein (50 nM final concentration) to the PBS-BSA solution. The microtiter plate was previously prepared with each cuvette containing 0.6 mM of S2302 diluted in PBS-BSA total volume of 90 µL, at precisely at 0.5, 1.5, 2.5 and 3.5 min, 10 μl of plasma-treated plasma kallikrein was collected and added to different cuvette wells. After 4.5 min, the microtiter plate was inserted into the spectrophotometer and absorbance was obtained at 405 nm for 15 min at room temperature. The half-life (T 1/2) of the kallikrein inhibition was determined by plotting the change in absorbance from 7 to 14 min and entering the ΔOD/min. The pseudo-first-order rate k’ constant was calculated by dividing the ΔOD/min into the natural log of 2, 0.693, manually or using Prism 10.0. To obtain a C1 inhibitor final activity level, the k’ constant was divided by the 2nd order rate constant of C1INH inhibition of plasma kallikrein 1.02 x 10^6^ M^-1^. Since the mouse plasma was not independently characterized for its concentration of C1INH, all final values were normalized by the international standard that 1 ml plasma contains 1 unit of plasma C1INH (73).

### Competitive ELISA for plasma high molecular weight kininogen (CELISA)

Plasma from children with CNS-M at hospital entry before artesunate treatment (CNS-M-En), CNS-M at discharge following clinical recovery (CNS-M-D), UM, acute fever non-malarial (AFN), and healthy community control children (Healthy) were obtained from whole blood collected into heparin anticoagulant. Plasma HK antigen levels in these samples were assayed by a competitive enzyme-linked immunosorbent assay (CELISA) as described previously using a human polyclonal anti-HK that detects both intact and cleaved HK (75,76). The standard curve for the HK CELISA was performed with total kininogen deficient plasma reconstituted with known concentrations of purified HK. The HK assay has an intra-assay coefficient of variation (CV) of 7.7% and 8.6% with pure HK in total kininogen deficient plasma and normal human plasma, respectively, and an inter-assay CV of 8.8% of 10 runs over a 1-month period.

### Immunoblotting

Normal human plasma, plasma from malaria patients, and infected and non-infected mice plasma were prepared for immunoblot for HK, PK, FXII, and C1 inhibitor by adding 5 μl and 10 μl of a 1:10 dilution of human or murine plasmas, respectively, in Laemmli sample buffer containing 5% β-mercaptoethanol and heated at 95°C for 5 min. The relative concentration of protein in plasma was determined by performing immunoblots on 10-100% normal mouse plasma (NMP) samples prepared identically to unknown murine samples. All bands of the protein of interest (NMP or *PbA*-infected plasmas) were compared to transferrin and band intensity was determined by ImageJ. Samples were run on either a reducing 7.5% polyacrylamide Tris-HCl Criterion gel (Bio-Rad) or a 10% polyacrylamide Tris-Glycine gel, as indicated. In certain experiments, the gels were stained with Coomassie blue. In other experiments gel electrophoresis was followed by transfer to Immobilon-P membranes (Millipore, Bedford, MA, USA). Protein bands were visualized with ECL substrate or Licor. Western blots were quantified by densitometric analysis (ImageJ). When immunoblots were used for quantitative analysis, transferrin or Ponceau S staining was used as loading controls.

Several antibodies to HK were used in these investigations. A goat polyclonal antibody to HK was used initially to immunoblot human HK in malaria patient plasmas to determine if there was cleaved HK and was used for the HK CELISA (see below) (76). This antibody detects intact and cHK. Cleaved HK was determined by the presence on immunoblot of the 56 and/or 46 kDa band(s) of the light chain of HK on samples reduced with 5% β-mercaptoethanol and heated for 5 min prior to addition to the SDS-PAGE for later transfer for immunoblot. A rabbit polyclonal antibody reared to the human HK D5 peptide LDDDLEHQGGHVLDHGHKHKHGHGHGKHKNKGKKNGK was also used to immunoblot human plasma HK. Two different rabbit polyclonal antibodies with similar sequences were prepared to detect murine HK D6 by immunoblot: anti-Cys-AHDDLIPDIHVQPDSL-SFKLISDFPEAT or - AHDDLIPDIHVQPDSLSFKLISDFPEATSPK. Finally, a second rabbit anti-mouse HK D5 antibody, anti-Cys-HGHGHGHGHGHGKHTNKDK, also was utilized for immunoblot investigations on mouse Cys-AHDDLIPDIHVQPDSLSFKLISDFPEAT and Cys-HGHGHGHGHGHGKH-TNKDK were prepared at BIOMATIK (Kitchener, ON, Canada). The other antibody was generously provided by Dr. Keith McCrae, Cleveland Clinic Foundation. In certain experiments, the relative antigen concentrations of HK, PK, FXII, and C1 inhibitor in plasma from murine *P. berghei* ANKA infected blood were estimated by scanning the intensity of the antigen band(s) on immunoblots. Each sample’s antigen level was compared to at least 9 normal mouse samples run on the same immunoblots after the values were normalized for loading with transferrin or Ponceau S. Note: when using an anti-mouse D6 antibody, two bands for HK are present. Quantitative studies include both bands in the amount present in normal and *PbA*-infected murine samples. Results are expressed at IU/ml antigen concentration, 1 unit being the amount present in 1 ml of pooled normal mouse plasma.

### Bradykinin assay

Blood samples from uninfected and *PbA*-infected mice for the BK assay were collected into a cocktail of 7 protease inhibitors diluted 1:9 with blood at the final concentrations indicated: hexadimethrine bromide (3 mg/ml), nafamostat mesylate (6.9 μg/ml), EDTA (6.1 mM), sodium citrate (12.4 mM), formic acid (1%), omapatrilate (1 μM) and chloroquine (1 mM) (23). Plasma was prepared by centrifugation at 1200 xg for 5 min at room temperature in an Eppendorf centrifuge. The collected plasma was frozen at-80°C until assay. Standards for the assay was the addition of known quantities of bradykinin to murine HK deficient plasma from *Kng1^-/-^* mice whose plasma contains the same cocktail of inhibitors. The assay was performed at the Lerner Research Institute, Mass Spectrometry Laboratory for Protein Sequencing at the Cleveland Clinic, Cleveland, OH. At the time of assay, 100 μl samples were processed using a Waters Oasis μElute 96-well plate according to the manufacturer’s recommendations. The eluted samples were dried in a Speedvac and reconstituted in 40 μl 0.1% formic acid. The LC-MS/MS (ThermoFisher Altis plus triple quadrupole mass spectrometer running in positive ion mode) was performed by injecting 20 μL of each processed sample onto a Gemini® 3 μm C18 110Å LC 150 x 2 mm column using a Vanquish UHPLC running at 0.30 mL/min with 3.2% DMSO in water and 0.1% formic acid as solvent A and 3.2% DMSO in methanol and 0.1% formic acid as solvent B. The data were processed using Thermo XCalibur software package. A processing method was developed using the transitions and calibration curve made by the standard samples, and the results were generated in Thermo XCalibur Quant Browser.

## Statistical Analysis

Data from the patient and ECM studies were compared in the text and graphed as the mean ± SD with individual values indicated unless otherwise stated. Each point represented an individual patient or animal. The total number of patients or animals in each group was described in figure legends. All data of various animal groups were examined for Gaussian (normal) or nonparametric distribution (e.g., Shapiro-Wilk). For data that were normal in distribution, comparison between 2 groups was performed by a two-tailed t-test and 3 groups by ANOVA. In human studies, nonparametric data was evaluated by Mann-Whitney or Kolmogorov-Smirnov tests. An unpaired t test was used to determine differences between groups. Statistical significance was defined as a P < 0.05. Kaplan-Meier curves were used to characterize survival curves. The Mantel-Cox and Gehan-Breslow-Wilcoxon tests were used to determine significant differences. The plasma biomarkers were nonparametric in distribution. Pearson correlation was used to assess the relationship between HK levels and biomarkers of vascular activation, cytokines, and coagulation/fibrinolysis. Statistical data were analyzed using PRISM 10.6.1

### Study Approvals

Ethical review and approval for the collection of Kenyan children’s samples was obtained from the Kenya Medical Research Institute (KEMRI) Scientific and Ethical Review Unit (SERU Protocol Number 3485) and University Hospitals Cleveland Medical Center/Case Western Reserve University (CWRU) Institutional Review Board (Number 12-16-15). Approval for collection of human plasma from normal human donors for assay standardization at CWRU was IRB CASE 19Z05, Number 01-06-02.

All studies were approved by Case Western Reserve University Institutional Care and Use Committee (IACUC) (Protocol #2014-0089) and performed in accordance with the guidelines of the American Association for Accreditation of Laboratory Animal Care and the National Institutes of Health.

Experiments were performed using 6-to 8-week-old male and female mice.

## Data availability

All supporting data values provided in the figures, text, and supplement is submitted in one Excel file. Each tab refers to all the data in each panel of each figure in the manuscript text and Supplement. All reagents and animals used in these studies are available upon request to the authors.

## Acknowledgements

We would like to acknowledge Drs. Markus Schwaninger and Walter Hauser from the University of Lubeck for generously providing the Slco1c1-CreERT2 mice that selectively targets brain vessel endothelium. Also, we want to thank the contributions of Drs. Ling Li and Belinda Willard at the Metabolomic and Proteomic Laboratory, Lerner Research Institute at the Cleveland Clinic Foundation for their assistance is establishing the mass spectrometry bradykinin assay. Last, we want to thank Dr. Jeffrey Breit of Rezolute, Inc. for providing the plasma kallikrein inhibitor for these studies.

## Author contributions

AHS, JWK, and AASP conceived the project. ASP, DET, YJS, AAM, SS, RPSA, YSP, TG, BBB, and AED performed critical research. JWK, DM, SO, YSP and AED acquired the clinical patient samples. AASP, JWK, and AHS wrote the manuscript. JS, CCN, OM, DG, MB, PJR, CJJ, KRM, AAPCN, BBB and AED edited the manuscript. AHS is responsible for the final version.

